# Aging and reorganization of the neural hierarchy underlying action perception: older adults prioritize affective-communicative information

**DOI:** 10.64898/2026.08.18.745087

**Authors:** Xue-Rui Peng, Qiaoli Huang, Angelika Lingnau, Christian F. Doeller, Shu-Chen Li

## Abstract

Everyday action perception unfolds in complex contexts, requiring coordinated processing from low-level visual information to higher-level inferences about action types and social interactions. Aging is accompanied by sensory and cognitive changes, raising the question of how these processes may be altered. Here, we combined action similarity judgments with analyses of time-resolved and source-localized brain activity measured with magnetoencephalography (MEG) to investigate age differences in action perception when younger and older adults viewed naturalistic actions. Using cross-validated variance partitioning within a multivariate representational framework, three key findings were uncovered. First, affective-communicative features predicted similarity judgments better than, and independent of, visual and action-related features in both age groups. Second, neural processes in both groups followed a hierarchy from visual to action-related and affective-communicative features and recruited similar functional regions. However, older adults showed delayed visual processing but earlier affective-communicative processing, indicating selective prioritization rather than uniform slowing of subprocesses. Third, when sensory uncertainty was increased through visual blurring, older adults showed hierarchical reorganization, with affective-communicative features emerging earlier than action-related features, whereas younger adults preserved the original hierarchy. Together, these findings unveil adapted information prioritization in healthy aging, whereby socially meaningful information increasingly guides action perception when sensory input becomes unreliable.

## Introduction

Action perception—the ability to perceive and interpret observed actions of others—is fundamental to everyday functioning and social interaction ^1^. In daily life, actions rarely occur in isolation but unfold within rich contextual backgrounds. Perceiving and understanding such naturalistic actions require integrating information across multiple processing levels, from extracting basic visual features to identifying action categories and making higher-order inferences about affective states and communicative intents ^2–4^. Current evidence on neural processes of action perception reveals a sequence of spatiotemporal processing in younger adults ^2,4,5^. Regarding brain regions, visual features, such as the gist of the scenes (e.g., spatial scale and structure), are processed in occipitotemporal visual regions ^4^. Processing action-related features and categories primarily involves the lateral occipitotemporal and parietal cortices that constitute the core action observation network ^5–8^. Social and affective features, such as communicative exchanges of the observed actors, are processed in a distributed brain network including the insula, amygdala, and superior temporal sulcus, extending beyond the action perception network to support social and affective processing ^3–5,9,10^. In terms of dynamics, recent EEG studies showed that processes across these levels follow a temporal hierarchy from visual, over action, to social-affective features in younger adults, with the processing of higher-level features contributing most to intuitive action understanding occurring later in time ^2,11^.

Even without pathological brain changes, healthy aging is accompanied by gradual declines in the speed and precision of sensory and cognitive processes ^12–15^, visuomotor integration efficiency ^16,17^, as well as mechanisms of perceptual and cognitive inference ^18,19^. Together, these changes may affect the underlying mechanisms of action perception in older adults ^20,21^. To date, how aging may influence the spatiotemporal processing of action perception is unclear, particularly concerning the potential interplay between neurocognitive declines and adapted alternative processing operations. On one hand, aging is characterized by sensory deficits and reduced processing fidelity ^12,13,22,23^. Age-related sensory declines have been linked to noisier neural information processing, particularly in occipital and temporal cortices, which reduces the signal-to-noise ratio of sensory encoding and compromises the fidelity of perceptual representations ^24,25^. Besides perception, aging is also associated with dedifferentiated representations of cognitive processes ^13,26^ and the coupling between cognition and sensory functions ^12,14^, which has been suggested to be, in part, attributable to reduced neuromodulatory regulation of neural processing fidelity, particularly due to age-related declines in dopaminergic modulation ^27–29^. On the other hand, other studies suggest that older adults may instead increase the engagement of adapted processing, such as a greater reliance on prior knowledge and context to guide perception ^20,21,30^, to deal with the less reliable sensory inputs. Similarly, whereas selective attention for suppressing irrelevant information is known to decline during aging ^31^, the socioemotional selectivity theory ^32^ posits a motivational shift towards focusing on socially and emotionally meaningful goals in older adults. This suggests a possible reallocation of cognitive resources to prioritize specific types of information for adapted processes ^33^.

However, it remains unknown how these factors may interact to influence action perception in older adults. To what extent are processes in the spatiotemporal hierarchy of action perception, as revealed in younger adults, affected by sensory and cognitive aging? Given age-related differences in motivational orientations, would older adults prioritize higher-level (e.g., affective-communicative) features during action perception more than younger adults?

### Study overview

To address these questions, we combined behavioral similarity judgments with magnetoencephalography (MEG) to characterize cortical temporal, spatial, and functional organization of action perception in younger and older adults to understand adult development of action perception (see Fig. 1 for an overview of the study design). The stimulus set comprised context-rich images depicting 16 categories of everyday actions varying in social communicativeness and emotional valence. In a pre-study (Fig. 1b), an independent sample of younger and older adults provided ratings of action-related and affective-communicative features of the stimuli. The main study consisted of two separate experiments with distinct groups of younger and healthy older adults. In Experiment 1 (Fig. 1c), participants arranged action images based on subjective perceived similarity of the actions. In Experiment 2 (Fig. 1d), participants underwent MEG recording while viewing the same set of stimuli as in Experiment 1 and the pre-study. To ensure that participants attended to all the stimuli during the experiment, they also performed a one-back action-category detection task. Critically, Experiment 2 included a within-participant manipulation of sensory uncertainty. We applied Gaussian blurring to simulate age-related decline in visual acuity ^22^. This manipulation enabled us to test how action-perception processes adapt when visual inputs are degraded. Original and blurred versions of each stimulus were presented in an intermixed, randomized order. To comprehensively characterize the nature of action perception in both age groups, we performed representational similarity analyses (RSA) with variance partitioning to examine age differences in cortical spatiotemporal processes of action perception. We hypothesized that: (i) the processing of affective-communicative features would emerge later but contribute more to intuitive action perception than lower-level features in both age groups; (ii) older adults may prioritize higher-level affective-communicative information given age-related sensory decline and the shift in motivational focus; and (iii) stimulus blurring would delay visual feature extraction, with potential downstream effects on higher-level processing in both age groups. To preview, we found that affective-communicative features dominated intuitive action perception both in younger and older adults, explaining most of the variance in behavioral similarity judgments of perceived actions. In both age groups, whereas cortical processes follow a temporal hierarchy from visual to action-related and affective-communicative features, older adults showed a prioritization of affective-communicative features, a pattern that becomes even more pronounced when visual inputs are degraded. These results reveal a reorganization of processes underlying action perception in old age that is aligned with the postulated motivational shift. Socially meaningful information is preferentially processed by older adults to guide action perception, particularly when sensory noise is high.

**Fig. 1.**
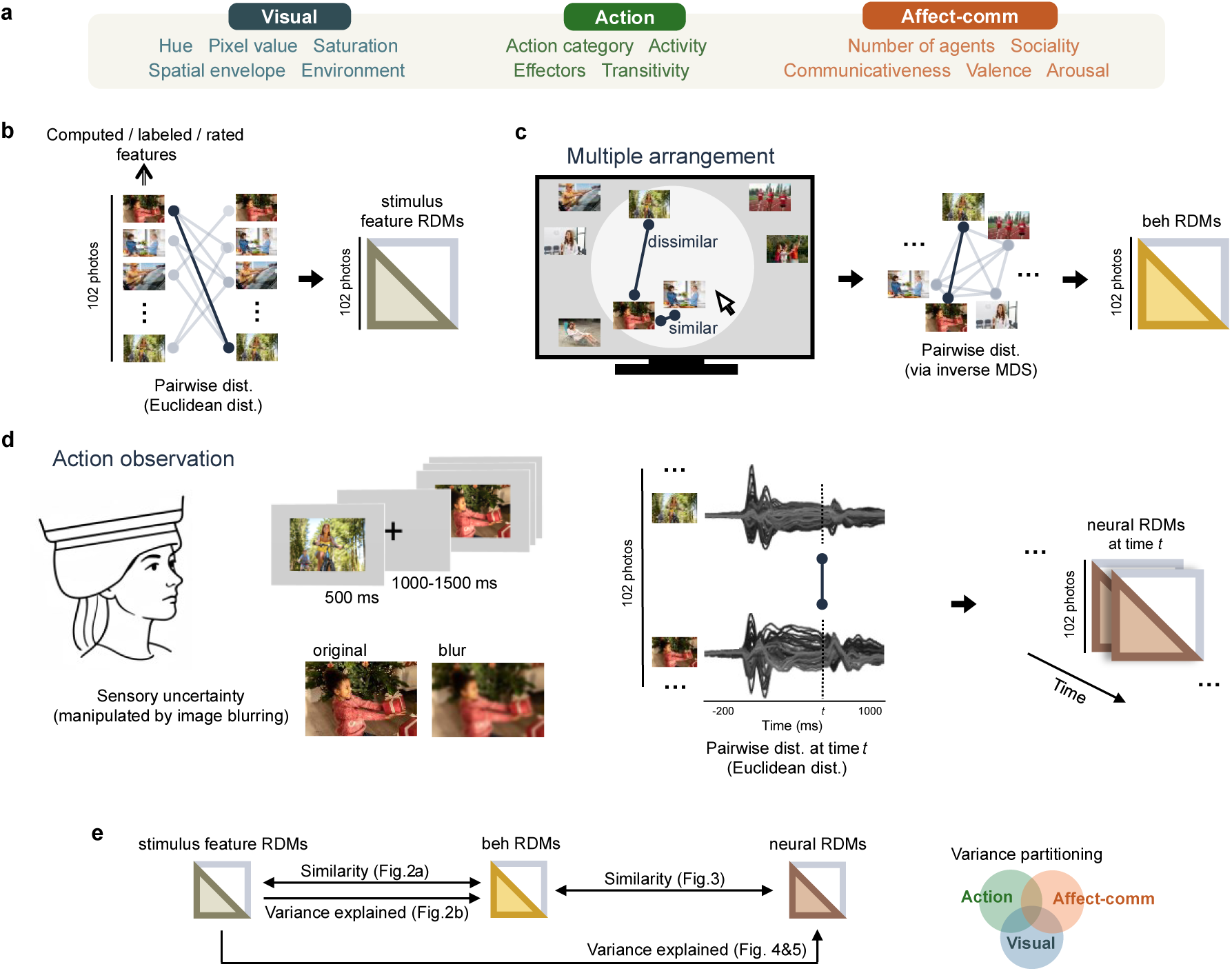
Study overview of experimental and analytic approaches for assessing age differences in action perception. **a** Stimulus features were obtained via algorithmic extraction (saturation, hue, pixel value, spatial envelope [GIST model]), experimenter annotations (environment, action category, effectors, number of agents), and behavioral ratings (transitivity, activity, communicativeness, valence, arousal, sociality) collected in the pre-study by an independent sample of younger and older participants (see Supplementary Text S1 and Fig. S1 for details of the pre-study). We grouped these features into feature sets (i.e., visual, action, and affective-communicative [affect-comm]). **b** Pairwise Euclidean distances were computed separately for each algorithmic, annotated, and rated feature across all stimuli (n = 102) to construct stimulus feature-specific representational dissimilarity matrices (RDMs). **c** In Experiment 1, participants arranged all stimuli by their perceived similarity in a multiple-arrangement task. Behavioral RDMs were derived from the resulting stimulus pairwise distances using inverse multidimensional scaling (MDS). **d** In Experiment 2, participants viewed the same set of stimuli during MEG recording, with sensory uncertainty varied by showing each stimulus in either its original or blurred version. For each condition, pairwise Euclidean distances across the 102 stimuli were computed at every time point of the MEG data to construct time-resolved neural RDMs. **e** Representational similarity analytic framework connecting stimulus features, behavioral judgments, and neural activity. Together, the analyses (1) assessed similarity by correlating behavioral RDMs with feature RDMs (results shown in Fig. 2) and time-resolved neural RDMs (results shown in Fig. 3), as well as (2) performed cross-validated variance partitioning to quantify the variance in behavioral (results shown in Fig. 2) and neural activity patterns (results shown in Figs. 4 & 5) explained by visual, action, and affective-communicative features. See Methods for details of the experimental tasks and analytic procedures. All images shown in this figure were licensed from Shutterstock (https://www.shutterstock.com/) and are reproduced with permission.

## Results

### Affective-communicative features explain most unique variance in intuitive action perception

To characterize how people perceive and categorize observed daily actions performed by others, we utilized a computerized multiple action arrangement task ^2,34^ in which participants were asked to place images depicting similar action closer together and dissimilar ones farther apart in a circular arena based on their own subjective judgments of similarity (Fig. 1c).

Behavioral representational dissimilarity matrices (RDMs) were constructed using inverse multidimensional scaling ^34^ from individual multi-arrangement data. Cross-participant consistency was quantified using leave-one-out correlations of the RDMs and exceeded chance levels in both age groups (Kendall’s *τ_a_* = 0.21 ± 0.13 for younger and 0.11 ± 0.05 for older adults; *ps* < 0.001). To identify which stimulus features drive perceived action similarity judgments, we correlated 14 candidate feature RDMs—spanning visual, action-related, and affective-communicative dimensions—with each participant’s behavioral RDM. These correlations were used for feature selection to define a common feature space to enable quantitative comparisons between age groups in subsequent analyses. An alternative approach—performing feature selection separately for each age group and taking the union of selected features—yielded the same set of features, confirming the robustness of our selection (see Supplementary Fig. S2 for age-specific analyses). Among the visual features, only environment—indicating whether an action occurred indoor or outdoor—contributed significantly to the similarity judgments. Several action-related features (action category, effectors, transitivity, and activity) and affective-communicative features (communicativeness, valence, and arousal) were all significantly correlated with the similarity judgments assessed by the action arrangement task (Fig. 2a). While intercorrelations among the stimulus features were strongest within each feature set (visual, action, affective-communicative), there were also some modest correlations across sets (spatial envelope [GIST-based descriptor] and effectors *τ_a_* = 0.14, spatial envelope and activity *τ_a_* = 0.14, effectors and communicativeness *τ_a_* = 0.14; see Supplementary Fig. S1c). Additional analyses were conducted to examine cross-participant consistency for features in the affective-communicative set and compare age effects at the individual feature level (see Supplementary Text S1 and Fig. S1a, S1b).

**Fig. 2.**
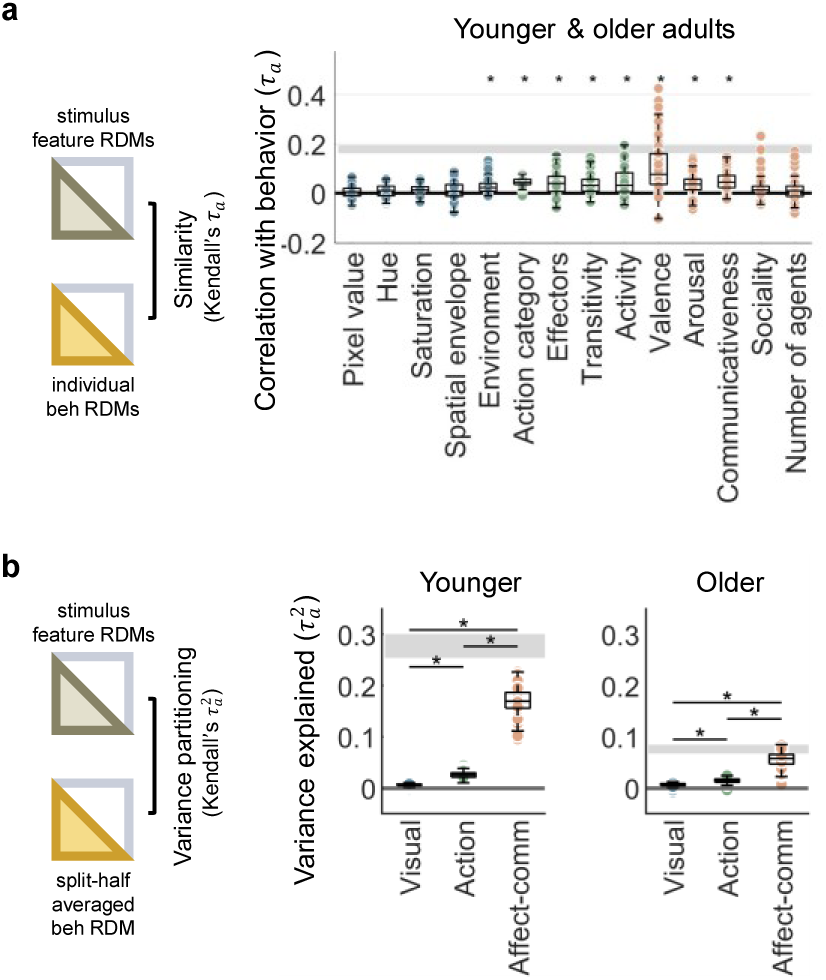
Stimulus feature contributions to perceived action similarity. **a** Correlations between participants’ behavioral representational dissimilarity matrices (RDMs) and stimulus feature RDMs in the entire sample of younger and older adults (Experiment 1). Each dot represents one participant (values of individual participants overlap). The noise ceiling is shown in gray. Asterisks indicate significant correlations (FWE-corrected *p* < 0.05, sign-permutation tests). **b** The unique variance explained by visual, action, and affective-communicative features separately for younger and older adults. Each dot represents one iteration from 100 split-half cross-validations. The split-half reliability of the data is shown in gray. Asterisks indicate significant differences between feature sets (all *p* < 0.001, Wilcoxon signed-rank tests). Affective-communicative features uniquely explain more variance in behavioral RDMs than visual and action features in both age groups.

To quantify the specific contribution of each of the three feature sets comprised of significant features identified above, we conducted split-half cross-validated variance partitioning analyses on data from Experiment 1 separately for younger and older adults to examine age effects (see Methods). The results revealed similarities but also differences between age groups. Given individual differences in action similarity judgments, the explainable amount of variance as indicated by the true split-half squared correlations were 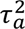 = 0.28 ± 0.02 and 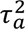 = 0.076 ± 0.01 for younger and older adults, respectively. The lower squared correlation in older adults reflects, in part, a lower cross-participant consistency of RDMs in older compared to younger adults (Mann-Whitney *z* = 3.79, *p* < 0.001).

Age differences in the amount of explainable variance notwithstanding, in general the affective-communicative features consistently contributed significantly more unique variance than visual or action features in each age group (all Wilcoxon *z* > 8.33, *p* < 0.001; Fig. 2b). In younger adults, the affective-communicative feature set contributed a significant amount of variance (*p* < 0.001, one-tailed sign permutation testing) and the action-related feature set reached marginal significance (*p* = 0.07), while the visual feature set did not (*p* = 0.37). The variance uniquely explained by affective-communicative features (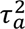 = 0.17 ± 0.02) accounted for 60% of the explainable variance. In older adults, both action-related and affective-communicative feature sets contributed significant amounts of variance (*p* = 0.004 and *p* < 0.001, respectively), while the visual feature set did not account for a significant amount of variance (*p* = 0.13). In older adults, the variance uniquely explained by the affective-communicative feature set (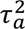 = 0.056 ± 0.01) amounted to 73% of the explainable variance. Further analyses revealed that valence contributed the largest amount of variance among the affective-communicative feature set in both age groups (see Supplementary Fig. S3), suggesting its key role in judging similarity of perceived action. Shared variance explained between feature sets was negligible for both age groups (all *p*s > 0.20, see Supplementary Fig. S4).

### MEG patterns are related to perceived action similarity in younger and older participants

To investigate the cortical processes of action perception, we conducted a MEG study (Experiment 2). To this aim, another sample of participants simply viewed the same 102 stimuli shown in Experiment 1 either in the original or blurred viewing condition, while performing a one-back action category detection task to maintain attention (see Methods). Note that the task accuracy did not differ significantly between younger (76.2% ± 10.5%) and older adults (71.4% ± 8.3%, *t*_61_ = 1.96, *p* = 0.054) nor between viewing conditions (*p*s > 0.16). Reaction time also did not differ between age groups (younger adults: 800 ± 127 ms; older adults: 799 ± 128 ms; *p* = 0.98). To examine whether brain activity patterns reflect perceived action similarity, we correlated the individual time-resolved neural RDMs (from Experiment 2) with a common aggregated behavioral RDM (across all participants from both age groups from Experiment 1). Since behavioral RDM reliability differs between younger and older adults, using a combined behavioral RDM provides a common basis for brain-behavior correlations that is not confounded by age differences in noise or consistency. Analyses using age-specific behavioral RDMs yielded comparable results (see Supplementary Fig. S5). In both age groups and conditions, this aggregated behavioral RDM correlated significantly with neural RDMs beginning shortly (around 120-140 ms) after stimulus onset and extending over a broad time window (Fig. 3). Pairwise cluster-based permutation tests revealed that the two age groups differed in how viewing condition affected brain-behavior correlations. In younger adults, no significant clusters revealed differences between the original and blurred conditions. By contrast, in older adults, there were two significant clusters (324–430 ms and 506–597 ms after stimulus onset) where the brain-behavior correlations are stronger in the original condition than in the blurred condition.

**Fig. 3.**
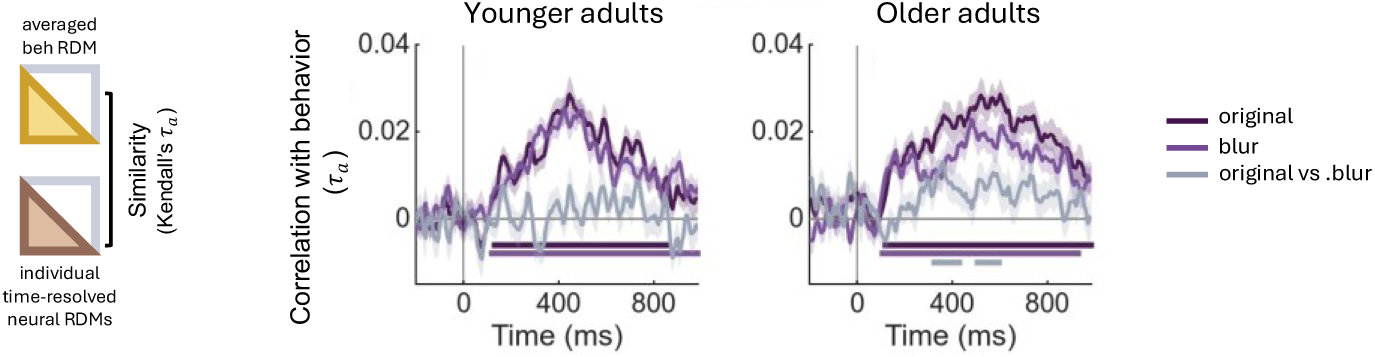
Neural activity tracks perceived action similarity over time. Time course of correlations between behavioral RDM (averaged across all participants from both age groups in Experiment 1) and individual neural RDMs (from Experiment 2) are shown for the two age groups and viewing conditions. Additional analyses using age-group specific behavioral RDMs were also conducted, which yielded similar results (see Supplementary Fig. S5). Significant time periods (sign-permutation testing, cluster-corrected *p* < 0.05) are indicated with horizontal lines below the curves (along the x-axis).

We next examined whether the stimulus features of the three sets (visual, action, and affective-communicative) that contributed to perceived action similarity judgments are also associated with MEG activity patterns. Specifically, we correlated the neural RDMs with each of the eight feature RDMs that were identified as significant predictors of action similarity judgments in Experiment 1 (see Fig. 2a). Across both age groups and viewing conditions, all feature RDMs showed significant correlations with neural RDMs (see Supplementary Fig. S6). These results suggest that the brain spontaneously processes these stimulus features during naturalistic action viewing even though participants in Experiment 2 simply viewed the actions during MEG scanning while performing a one-back task of action categories in a small number of catch trials.

### Common temporal hierarchy during action perception in younger and older adults

We next investigated the cortical temporal processing hierarchy of the feature sets contributing to action perception. Specifically, we identified the time windows in which the feature sets uniquely accounted for significant amounts of variance in brain activity (Fig. 4). When viewing original images, in younger adults, visual features dominated early processing stages (62–375 ms after stimuli onset), followed by action-related features in the mid-latency periods (203–627 ms), and affective-communicative features in an overlapping time window with action features but with a slightly later onset (264–662 ms). In older adults, a similar temporal progression across these three feature sets was also observed during the original viewing condition, with visual features accounting for unique variance in early time windows (67–425 ms), action features during mid-latency periods (234–329 ms, 345–602 ms), and affective-communicative features in a slightly later but overlapping period (329–793 ms, Fig. 4b). Because behavioral analyses indicated negligible shared variance among feature sets, we focus here only on the unique contributions (see Supplementary Fig. S7 for the results of corresponding analyses of shared variances).

**Fig. 4.**
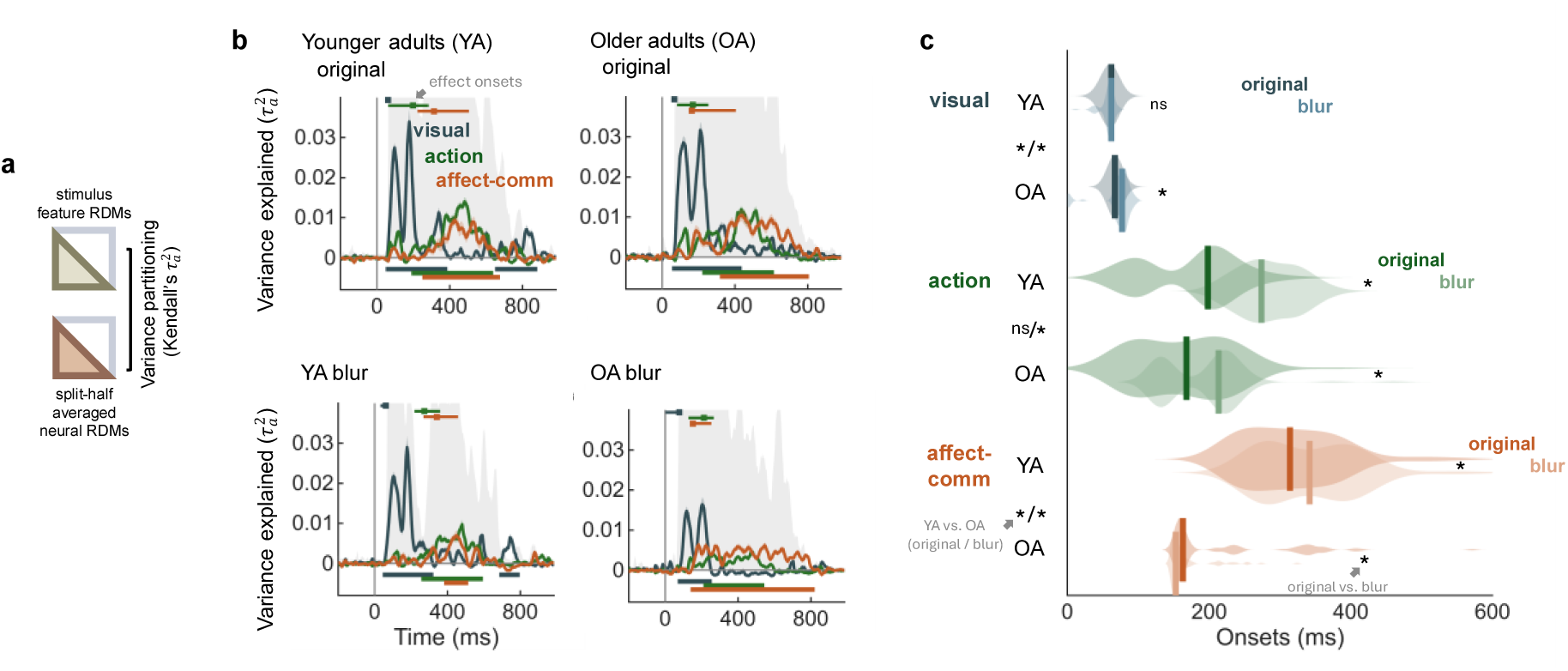
Brain dynamics uniquely associated with the processing of visual, action and affective-communicative features in two age groups and viewing conditions. **a** In each iteration, participants were randomly divided into two halves. Neural RDMs averaged from one half of the participants were used to fit full and reduced regression models, and model predictions were then correlated (Kendall’s *τ_a_*) with the averaged neural RDM from the held-out half of the participants to estimate the unique contributions of visual, action, and affective-communicative features (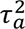). **b** Time courses of unique variance explained by visual, action, and affective-communicative feature sets for younger adults (YA) and older adults (OA) under the original and blurred viewing conditions. The subset of the feature RDMs were derived from subjective ratings (transitivity, activity, communicativeness, valence, and arousal) pooled across younger and older adults from a pre-study. Control analyses using age-group specific ratings also showed similar results (see Supplementary Fig. S8). Shaded areas show standard error; gray shaded backgrounds show the split-half reliability of the data. Significant time points (sign-permutation testing, cluster-corrected *p* < 0.05) are indicated with horizontal lines below the curves (along the x-axis). Horizontal bars at the top of each panel indicate the 95% confidence interval of effect onset for each feature set, with the dot marking the median onsets. Onset was defined as the starting time point of the earliest significant cluster (sign-permutation testing, cluster-corrected *p* < 0.05). **c** Distributions of effect onset for each feature set in the two age groups and viewing conditions across 100 split-half iterations. Vertical line indicates median onsets. Symbols between YA and OA labels along y-axis show age comparisons (left: original; right: blur); symbols next to distributions show viewing condition comparisons (original vs. blurred) within age groups. \**p* < 0.05; ns, not significant.

Onset analyses across 100 split-half iterations also confirmed the above temporal hierarchy in both age groups, with brain activities unfolding sequentially from visual to action and affective-communicative features (Fig. 4c). In younger adults, visual features emerged earliest (median onset = 62 ms, 95% CI: 47–67 ms), followed by action features (median = 198 ms, 95% CI: 62–284 ms), and finally affective-communicative features (median = 314 ms, 95% CI: 224–506 ms), with the later feature sets emerging significantly later than the previous one (all pairwise Wilcoxon signed-rank tests *z* > 8.34, *p* < 0.001). Brain activities of older adults showed a similar sequence, with visual features appearing at a median of 67 ms (95% CI: 62–77 ms), followed by action (median = 168 ms; 95% CI: 82–254 ms) and affective-communicative features (median = 163 ms; 95% CI: 153–405 ms). Although the group-level medians for the latter two feature sets were numerically close due to distributional skew, pairwise analyses confirmed the same hierarchical organization as in younger adults (all pairwise *z* > 4.30, *p* < 0.002). Albeit the comparable temporal progression across the three feature sets between age groups, age differences in onset latencies were observed. Compared to younger adults, older adults showed significantly delayed onsets for visual features (difference in medians: Δ = +5 ms, Mann-Whitney U test *z* = −10.37, *p* < 0.001), whereas affective-communicative feature onsets were significantly earlier than those of younger adults (Δ = −151 ms, *z* = 7.65, *p* < 0.001), and the onsets of action features did not differ between age groups (*p* = 0.24).

### Age-specific effects of sensory uncertainty on temporal hierarchy

To examine how sensory uncertainty caused by the blurred viewing condition may affect processes of action perception differently between younger and older adults, we performed the same variance partitioning analysis on brain activities in the blurred viewing condition as done above for data from the original viewing condition. As shown in Fig. 4b, in younger adults, the temporal pattern resembled that of the original condition, with visual features contributing unique variance in early time window (57–309 ms), action features during mid-latency period (269–582 ms), and affective-communicative features in a relatively narrow and late period (395–501 ms). Interestingly, in older adults a reorganization of this hierarchy was observed. While visual features still emerged early (82–244 ms), affective-communicative features (153–808 ms) spanned an extended interval that began prior to and encompassed the action feature window (244–531 ms).

Analyses of onset latencies for the feature sets lent further support for the results above and revealed a preserved sequence hierarchy in younger adults under sensory uncertainty, with visual features emerging earliest (median = 62 ms, 95% CI: 32–72 ms), followed by action features (median = 274 ms, 95% CI: 218–360 ms), and finally affective-communicative features (median = 342 ms, 95% CI: 268–458 ms; all pairwise Wilcoxon signed-rank test *z* > 4.89, *p* < 0.001). However, in older adults a different temporal hierarchy was observed with respect to onset latencies: while the onset of visual features still emerged first (median = 77 ms, 95% CI: 2–92 ms), affective-communicative features started earlier (median = 153 ms, 95% CI: 148–254 ms) than action features (median = 213 ms, 95% CI: 128–267 ms; all pairwise |*z|* > 4.08, *p* < 0.001), reversing the previously reported progression of action-related then affective-communicative features previously observed in younger adults ^2^ and in older adults under the original unblurred viewing conditions as reported above (Fig. 4c).

Permutation-based tests showed significant group × condition interactions for all three feature sets (all *p*s < 0.001), indicating that blurring affected processing onsets differently between age groups. A significant main effect of condition was observed for each feature set (all *p*s < 0.05). The main effect of group was significant for visual features (*p* < 0.001) and affective-communicative features (*p* < 0.001), but not for action-related features (*p* = 0.59). To further examine the age-specific effects of sensory uncertainty, we conducted post-hoc test that compare the onsets within each age group.

In younger adults, blurring did not affect the onset of visual features (Wilcoxon signed-rank test *p* = 0.14) but resulted in delayed onsets for action-related features (Δ = +106 ms; *z* = −8.40, *p* < 0.001) and affective-communicative features (Δ = +28 ms; *z* = −3.09, *p* = 0.002). In older adults, blurring slightly delayed visual (Δ = +10 ms; *z* = −4.70, *p* < 0.001) and action features (Δ = +30 ms; *z* = −6.18, *p* < 0.001) but resulted in slightly earlier onsets for affective-communicative features (Δ = −10 ms, *z* = 7.65, *p* < 0.001), suggesting a prioritization of affective-communicative information in the blurred condition.

### Spatial organization of action perception

To examine the cortical sources underlying the unique contribution of visual, action and affective-communicative features, we conducted cross-validated variance partitioning analyses on the source-localized MEG data separately for age groups and viewing conditions (see Fig. 5a). Statistical tests were first restricted to age-common time windows identified from the time-resolved results in periods covering significant effects observed both in younger and older adults (see also Methods). In younger adults, when viewing the original stimuli, the spatial distributions of variances of brain activity in the age-common time windows uniquely explained by the three feature sets were as follows: visual features were maximal in the early visual cortex (EVC), whereas action-related features were primarily localized to the lateral occipitotemporal cortex (LOTC, including the extrastriate body area, EBA) and the affective-communicative features were most prominent in the insula, with distributed contributions from posterior superior temporal sulcus (pSTS), posterior parietal (precuneus and parieto-occipital), and cingulate cortex. Similarly, in older adults, visual features uniquely explained maximal variance of brain activity in EVC, and action-related features in LOTC. The affective-communicative features uniquely explained maximal variance of brain activity in the temporoparietal junction (TPJ), extending into superior temporal, anterior inferior parietal lobe (aIPL), precuneus, and posterior cingulate regions.

**Fig. 5.**
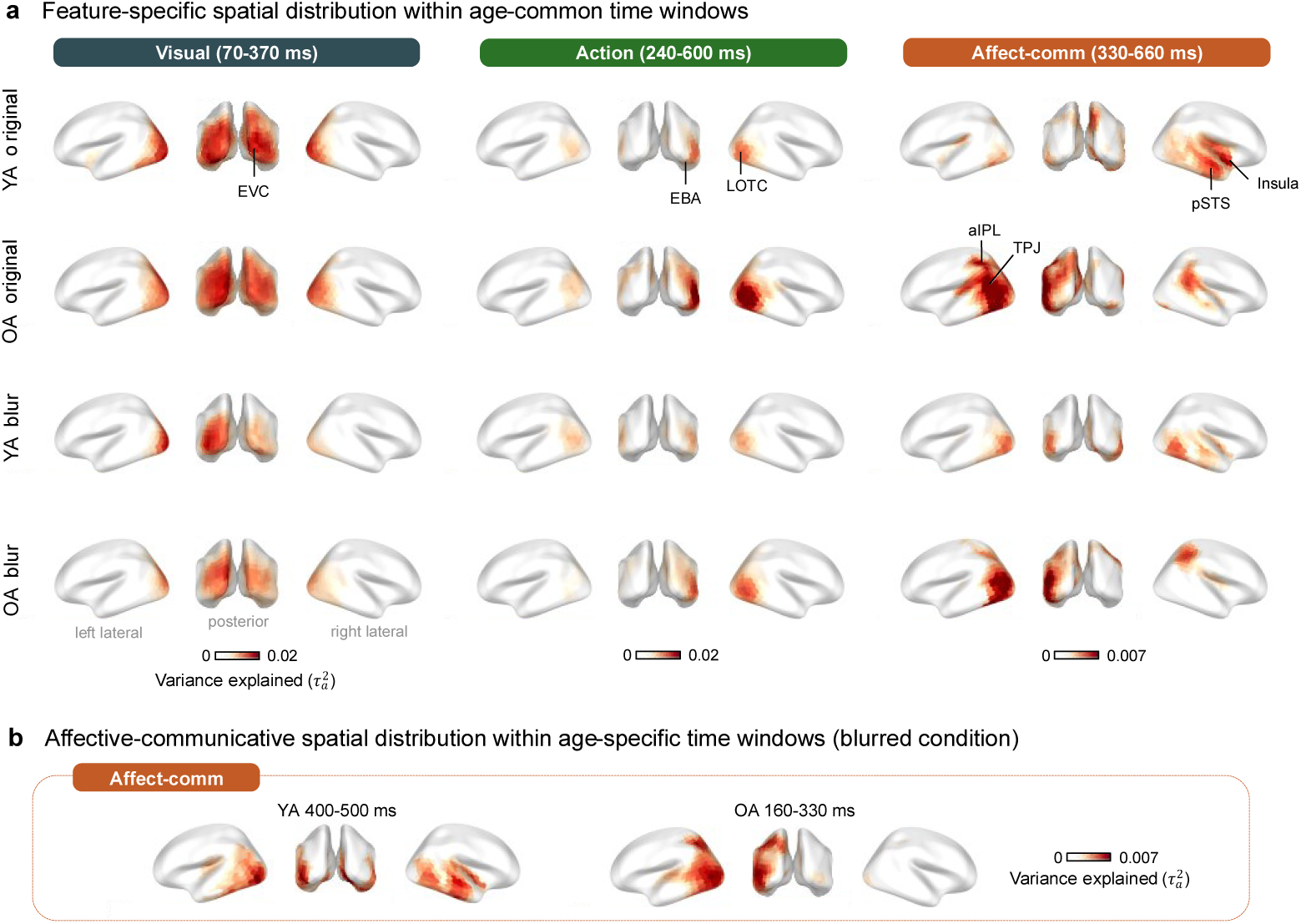
Cortical sources of variance in MEG activity uniquely explained by visual, action-related, and affective-communicative features. **a** Spatial distribution of unique variance in the two age groups (YA: younger adults; OA: older adults) and viewing conditions (original, blur). The age-common time windows for this analysis were determined by the overlapping significant time periods observed in the original viewing condition in both age groups (as shown in Fig. 4). **b** Spatial distribution of variance uniquely explained by affective-communicative features in the blurred condition during age-specific significant time windows. Only significant vertices were shown (permutation tests with 10,000 permutations, FWE *p* < 0.05). EVC, early visual cortex; EBA, extrastriate body area; LOTC, lateral occipitotemporal cortex; pSTS, posterior superior temporal sulcus; aIPL, anterior inferior parietal lobe; TPJ, temporoparietal junction.

In the blurred condition, the overall spatial pattern of unique contributions for visual and action-related features remained largely comparable to the original viewing condition in both age groups, with peaks in EVC for visual features and LOTC for action-related features. For affective-communicative features, during the age-common time windows, younger adults showed peaks in the posterior temporal cortex, with contributions from lateral occipital and ventral occipitotemporal regions. Older adults showed peaks in LOTC with distributed contributions from lateral occipital, posterior temporal, and posterior parietal cortex. Notably, given that the time course of affective-communicative processing in the blurred viewing condition differed markedly between age groups, we thus also performed further analyses tailored to the temporal profile of each group (Fig. 5b). For younger adults, affective-communicative information emerged later and within a narrower time window when viewing blurred stimulus compared with the original. Restricting statistical tests to this specific significant time window (400–500 ms) revealed spatial patterns that resemble those of the original condition. For older adults, affective-communicative processing emerged substantially earlier under blurred viewing. Focusing on this specific early time window (160–330 ms), during which affective-communicative processing was significant only in older adults, revealed spatial locations comparable to those observed in the original viewing condition (e.g., aIPL and TPJ), but already engaged during this earlier period. Together, these results suggest that under sensory uncertainty, older adults recruited affective-communicative-related networks with spatial patterns comparable to those in the original viewing condition but already at an earlier time window.

## Discussion

By combining representational similarity analyses of intuitive action similarity judgments with time-resolved and source-localized MEG activity, this study investigated adult age differences in the organization of cognitive and brain processes underlying action perception. Three key findings emerged. First, in both younger and older adults, affective-communicative features uniquely accounted for the largest amount of variance in how naturalistic everyday actions are perceived and categorized, above and beyond low-level visual and action-related features. Second, when viewing original stimuli, the brain activity of both age groups showed a temporal processing sequence related first to visual then action-related and affective-communicative features, which is consistent with previous findings in younger adults ^2^ and extends this temporal dynamic to older populations. Third, although older adults showed delayed neural processing onset of visual features relative to younger adults, the onset of brain activity associated with affective-communicative processing was earlier in older than in younger adults. Furthermore, when the visual input was degraded by blurring, the prioritization of affective-communicative features became even more pronounced in older adults, such that the processing of these higher-order features emerged much earlier than action-related features. This revealed an adapted temporal process of action perception that differs from the typical processing sequence observed in younger adults and in older adults when no sensory challenge was imposed. Together, these findings reveal a reorganization of action perception during adult development, whereby older adults prioritize the processing of higher-order affective-communicative features which are most relevant for the perception of others’ action and social cognition.

Our behavioral results showed that affective-communicative features played a central role in action similarity judgment. In both age groups, affective-communicative features independently accounted for the variance in intuitive action similarity judgments more than visual and action-related features (Fig. 2), with valence emerging as the most influential single feature (Supplementary Fig. S3). This pattern closely aligns with recent studies in younger adults ^2,35^ and underscores the inherently social nature of human action perception ^36^. Relative to younger adults, cross-participant consistency of action similarity judgment is lower in older adults, which reflects more heterogeneous judgments in this age group. Nevertheless, affective-communicative features accounted for a substantial proportion of the explainable variance in older adults. The greater interindividual difference in behavioral judgments among older adults observed here is consistent with age-related increases in performance heterogeneity ^37^ and intraindividual variability ^13,29^. Taken together, results at the behavioral level from Experiment 1 suggest that the constituent subprocesses for processing visual, action-related and affective-communicative features are also all still operative during healthy aging, albeit the precision of perceptual representations declines.

Using time-resolved representational analyses of brain activity assessed with MEG from Experiment 2, we found that both younger and older adults showed a cortical temporal processing sequence of action perception that progresses from visual to action-related and affective-communicative features. This finding further suggests that the fundamental mechanisms of action perception remain preserved in older adults. Importantly, results from Experiment 2 also uncovered an adapted processing prioritizing higher-order affective-communicative features in older adults, particularly when sensory uncertainty is high due to degradation of stimulus inputs.

Even during healthy aging, sensory declines contribute to delayed visual processing at multiple levels, including slower information transmission along the early visual pathway ^38^, delayed accumulation of visual information ^39,40^, and postponed emergence of representations in high-level ventral visual cortex ^41^. Consistent with these findings, in the original viewing condition we observed delayed onsets of visual feature processing in older adults compared to younger adults. Notably, however, despite delayed visual processing, older adults showed comparable onset timing for processing action-related features and a substantially earlier onset (about −150 ms) for processing affective-communicative features compared to younger adults. These findings shed light on the question regarding how age-related sensory and cognitive changes affect the temporal hierarchy of action perception. Rather than a uniform slowing across all constituent processes of action perception, aging appears to involve an adapted processing operation that reprioritizes the subprocesses. Importantly, the earlier processing onset of affective-communicative features in older adults should not be interpreted as faster or more efficient processing overall, but rather as a shift in the temporal allocation of processing across feature sets.

Increasing sensory uncertainty by degrading visual input through blurring, we examined how sensory challenges may interact with age differences in motivational factors to impact subprocesses of action perception. We discovered a pronounced reorganization of the processing hierarchy specifically in older adults. Under blurred viewing conditions, affective-communicative features emerged earlier than action-related features in older adults, reversing the typical temporal order observed under original viewing conditions. In contrast, younger adults maintained the original hierarchical sequence, showing only delayed onsets for action-related and affective-communicative features without altering their relative ordering. This age-specific reorganization reflects an adapted processing operation in which older adults reprioritize processing first towards higher-order affective-communicative features, which play the key role in intuitive action organization, to guide the processing of the other two lower-level subprocesses when visual input is degraded. This finding is in line with earlier evidence showing the increased reliance of older adults on prior knowledge and internal predictions when sensory signals are compromised ^20,30^. During action prediction, for instance, older adults show reduced sensitivity to early kinematic cues but greater use of contextual information to guide their responses ^21^. Similar shifts toward greater reliance on acquired priors have also been observed in visual and multisensory perception ^42,43^, as well as in naturalistic visual search, where older adults rely more heavily on prior expectations to guide eye movements than younger adults ^44^.

At a different level, our findings also lend support and extend the socioemotional selectivity theory ^32,45^ of adult development and motivated cognition ^46^ to the domain of action perception. Specifically, this theory proposes that as perceived future time horizons shrink, individuals increasingly prioritize emotionally meaningful and socially relevant goals. While such motivational shifts have been shown to influence attention ^46,47^, memory ^48^, and decision-making ^49^ in later life, our findings suggest that these motivational changes may also shape perceptual processing dynamics during action perception.

Of note, even though our one-back action detection task during the catch trials required participants to focus on the action category instead of features in the affective-communicative set, older adults still prioritized the processing of affective-communicative features. The number of catch trials was only a small fraction of the entire task in Experiment 2; however, performing the one-back task could still have some influence on the experimental trials. In younger adults, the processing time window for affective-communicative features was narrower under blurred compared with original viewing conditions, which may reflect a processing bias influenced by the catch trials. In older adults, however, the processing of affective-communicative features in the blurred condition started earlier and remained prolonged even when such information was unnecessary for task performance in the catch trials. This pattern suggests that the prioritization of socially meaningful information in aging may operate at a relatively higher-order level, such as a motivational shift toward social and emotional goals that modulated further cognitive and perceptual processes.

Our source-localized analyses revealed both similarities and differences between younger and older adults in the brain regions that support action perception. In both age groups and viewing conditions, visual features peaked in EVC, consistent with previous findings that low-level image statistics and scene properties are mainly processed in early visual areas ^4^. Action-related features peaked in LOTC, a region widely reported for processing body-related information, action semantics, and perceptual components of observed actions ^5,7,50^. The stability of these spatial patterns across age groups and viewing conditions suggests that fundamental visual and action processing are preserved in healthy older adults and are relatively robust to perceptual degradation. The two age groups differed, however, in the cortical regions engaged during affective-communicative processing. In younger adults, affective-communicative features were primarily associated with regions involved in affective evaluation and social cue processing, particularly the insula and pSTS ^9,10,51,52^. In contrast, older adults showed affective-communicative processing mainly in regions implicated in social inference and the interpretation of communicative intent, such as TPJ and aIPL ^1,3^. This difference in spatial localization converges with our behavioral findings which show that although valence was the dominant contributor among affective-communicative feature set in both age groups, communicativeness emerged as an additional significant contributor specifically in older adults (see Supplementary Fig. S3). Under the blurred viewing condition, affective-communicative processing in younger adults was both delayed and temporally compressed. Within their narrowed time window (370–500 ms), the spatial pattern closely resembled that observed under original viewing condition, suggesting that the neural substrates for affective-communicative processing remain stable despite temporal delays caused by degraded sensory inputs in younger adults. In older adults, by contrast, cortical activations comparable to the original viewing condition were already evident within an early time window (160–330 ms), and even before affective-communicative effects emerged in younger adults. This early emergence of similar patterns of brain functional recruitment in older adults is consistent with predictive coding accounts proposing that accumulated experiences or other contextual goals provide stronger priors when sensory input is degraded ^42^.

Several limitations should be considered when interpreting our findings. First, our stimuli were static images. While this allowed for experimental control of temporal complexity, natural actions unfold dynamically over time. Motion information carries rich cues about action goals, emotional states, and social intentions ^53^. Future studies should examine whether the prioritization of affective-communicative features observed in older adults becomes even more pronounced with dynamic stimuli, where older adults might leverage their preserved ability to extract biological motion information ^54^ and motivational shift to socially relevant information ^45^ to compensate for reduced sensitivity to static visual details. Second, our sample comprised healthy, cognitively normal older adults (MoCA scores ≥ 23). The pattern of results might differ in pathological aging such as mild cognitive impairment or dementia, where both sensory processing and social cognition are often compromised ^55,56^. Examining whether the adapted prioritization strategy we observed here may break down or be preserved in older individuals experiencing neurodegeneration could provide insights into when compensatory mechanisms may fail and potentially identify other markers of pathological aging that go beyond cognitive decline. Third, while we manipulated sensory uncertainty through Gaussian blurring to simulate age-related visual decline, this represents only one dimension of sensory degradation. Age-related changes also affect contrast sensitivity, visual processing speed, and attentional selection ^57,58^. Future studies could examine how other forms of sensory challenge (e.g., reduced contrast, brief presentation, divided attention) affect the hierarchical organization of action perception.

Taken together, our study shows that healthy aging is associated with a reorganization of processes underlying action perception. While both younger and older adults organize observed actions primarily along the affective-communicative features, older adults show a temporal prioritization in affective-communicative processing that becomes even more pronounced under sensory uncertainty. These results also lend support and extend socioemotional selectivity theory to action perception, suggesting that motivational priorities in later life may influence the temporal dynamics of how older adults perceive others’ actions. More broadly, we show that cognitive aging involves not only decline but also strategic reorganization of processes such that the adapted processing may be more in line with higher-order goals or motivational orientations, with the aging brain adapting its processes to optimize extraction of information most relevant for navigating the social world or other task requirements given its available neurocognitive resources.

## Methods

Two experiments were conducted to characterize multidimensional representations of observed actions at the behavioral (Experiment 1) and brain (Experiment 2) levels as well as their relations (see Fig. 1 for an overview). Prior to the experiments, an online pre-study with 125 participants (62 younger adults and 63 older adults) who rated the action and affective-communicative features of the stimuli was conducted (see Text S1 and Fig. S1 in the Supplementary Materials for details about the pre-study). All experiments were approved by the ethics committee of the University of Leipzig (341/23-ek). Informed consent was obtained from all participants before participating in the study.

### Participants

#### Experiment 1: multi-arrangement task of images depicting actions

Forty-five younger and 45 older adults took part in Experiment 1. One younger adult was excluded due to not completing the experiment. The final sample includes 44 younger adults (age range 19-29 years, mean age ± SD: 23.91 ± 2.67 years; 24 females, 20 males) and 45 older adults (age range 65-86 years, mean age ± SD: 72.77 ± 4.95 years; 22 females, 23 males).

#### Experiment 2: MEG experiment of action observation

A separate sample of 32 younger and 32 older adults took part in Experiment 2. All participants were right-handed, had normal or corrected-to-normal vision, without neurological or psychiatric conditions. Older adults were screened for cognitive impairments using the Montreal Cognitive Assessment (MoCA; cutoff ≤ 23) ^59,60^. One older adult was excluded due to random bursts of high-frequency interference during the second half of the MEG data collection. The final sample consists of 32 younger adults (age range 20-31 years, mean age ± SD: 25.16 ± 3.63 years; 17 females, 15 males) and 31 older adults (age range 65-77 years, mean age ± SD: 71.58 ± 3.44 years; 14 females, 17 males; MoCA score mean ± SD: 27.48 ± 1.90). We also characterized the sample using the Spot-the-Word Test ^61^, Identical-Pictures Test ^62^, a visual search task (T among L distractors) ^63^, and the Duke Social Support Index ^64^. As expected for samples with healthy older adults, they showed higher verbal knowledge, but slower processing speed and visual search efficiency compared to younger adults. Social support and interaction were comparable across groups (see Supplementary Table S1 for more details).

#### Stimuli

Previous studies have shown that social features drive the perception of observed daily actions in younger adults ^2,4,65^. Going beyond prior studies, we improved the stimulus set by systematically varying the stimuli along feature sets. Specifically, we carefully selected eight daily communicative actions and eight non-communicative actions, while also balancing the emotional valence of the stimulus set. Furthermore, to minimize the association between the number of agents and social factors, each action was depicted in three single-agent and three multi-agent images. Altogether our stimulus set included 16 actions, with six images for each action. It is important to note that we labelled action category based on the most salient action depicted in the images, but the images are embedded in rich contexts and may involve other peripheral actions. The labels and the categories were made for study design and data coding purposes, participants were not informed/instructed about the categories or labels. For instance, in the “arguing” category, one image shows a woman sitting in the driver’s seat and arguing with someone on the street, thus this image also entails the action of “driving.” Moreover, to add variations in the stimulus set, we selected 6 control images that depict outdoor and indoor scenes that show no actions or agents in the images. These control images were rated along with the images depicting action in the pre-study and were included in all subsequent analyses (Experiments 1 and 2). In total, our stimulus set included 102 stimuli (i.e., 96 action images + 6 control images, see Supplementary Fig. S9 for an overview of the stimuli). All images used in the present study were obtained under license from Shutterstock (https://www.shutterstock.com/).

### Experimental procedures

#### Experiment 1: multi-arrangement task

We adapted the multi-arrangement task ^34^ (e.g., see Fig. 1c) implemented on the MEADOWS platform (https://meadows-research.com/). Participants arranged 102 action images within a circular arena (Fig. 1) by dragging and dropping them such that the on-screen distances reflected their perceived similarity. No explicit instructions about action categories, similarity, or criteria were provided on which features to focus on, allowing the participants to sort the images based on their subjective, perceived similarities (see Supplementary Materials for task instructions). To accommodate all items, stimuli were shown as thumbnails, and a larger preview appeared while participants hovered over it (see Supplementary Fig. S10 for a screenshot of the interface). Dissimilarity matrices were estimated using Inverse Multidimensional Scaling ^34^. After an initial arrangement of all 102 stimuli, subsequent trials adaptively resampled subsets with weakest dissimilarity evidence using the ’lift-the-weakest’ algorithm. The final RDM was estimated by a weighted average of iteratively scale-adjusted distances ^34^. The task continued until the evidence criterion of 0.5 was reached for each pair or after 120 minutes. The task took on average 107 ± 19 min (196 ± 60 trials) for younger and 118 ± 5 min (136 ± 63 trials) for older adults.

#### Experiment 2: MEG experiment of action observation

During the experiment, participants simply viewed the same images depicting daily actions used in Experiment 1. No explicit instructions about what constituted an action were given nor any sorting was involved. To maintain attention during the experiment while minimizing cognitive load, participants performed a modified one-back task during occasional catch trials: when the image was presented together with a prominent frame, they were instructed to press a response button if they thought the image shown in the current trial depicted the same action category as the image shown in the previous trial (60%) and withheld responses otherwise (40%). Each of the 96 action images served once as a catch trial in each viewing condition (192 catch trials total), distributed evenly across five blocks. Catch-trial sequences were counterbalanced across the two viewing conditions. Catch trials were excluded from analyses. While Experiment 1 presented images only in their original form, Experiment 2 included two viewing conditions: original and blurred images. The degree of blurring was determined based on a pilot study, with the aim of applying substantial blurring while ensuring that accuracy of the one-back task in the catch trials remained comparable to that of the original images. Blurring was implemented by convolving each 400 × 600 pixel image with a two-dimensional Gaussian kernel (σ = 5; effective filter size ≈ 21 × 21 pixels) using MATLAB’s imgaussfilt function, which reduced fine-grained visual detail while preserving the overall spatial layout and structure (see Supplementary Fig. S11 for example images before and after blurring). The two conditions were randomly intermixed. Within each block, the 102 images were shown four times (twice per viewing condition) in a pseudorandomized order with no consecutive repetitions. Altogether, the task comprised 2,232 trials (2,040 experimental and 192 catch trials). As shown in Fig. 1d, each trial started with the presentation of a black fixation cross on a light gray screen (jittered duration: 1000–1500 ms, uniform distribution), followed by stimulus presentation (500 ms). The average experiment duration was 73.84 ± 3.94 minutes. We provided self-paced breaks every 22 trials, with a minimum 15-second pause in the middle of each block.

#### MEG Data Acquisition

MEG data were acquired in a magnetically shielded room using a 306-channel MEGIN Vectorview system (204 gradiometers, 102 magnetometers). Before the MEG recording, we first attached five head-position indicator coils to the participant’s head (three on the forehead and one on each mastoid). We then used an electromagnetic digitizer to digitalize three fiducial points (nasion and left/right preauricular points) and the 5 HPI coils, followed by whole head shape digitization (> 500 points) to enable accurate co-registration with individual structural MRI scans to improve source localization. These structural images were retrieved from the MPI-CBS database for participants with existing data from prior studies or newly acquired for the current study if a structural image was not available. MEG data were continuously acquired at a sampling rate of 1000 Hz with an online low-pass filter with a cutoff frequency of 330 Hz.

#### Data analysis

Analysis of multi-arrangement task (Experiment 1)

To understand how different stimulus features (i.e., visual, action-related, and affective-communicative features) contribute to naturalistic action perception, we conducted representational similarity analysis (RSA ^66^). Fourteen stimulus features were quantified by algorithmic extraction, experimenter annotations or participant ratings from the separate online pre-study (see Supplementary text S1 and Fig. S1 for details). To generate feature representational dissimilarity matrices (RDMs), Euclidean distances were then computed between all stimulus pairs. Visual features included pixel value, hue, saturation and spatial envelope (GIST ^67^), and environmental settings (i.e., indoor or outdoor). Action-related features comprised participant ratings of transitivity and activity, and experimenter-coded action categories and effector involvement (face, hands, arms, legs, torso). Affective-communicative features included participant ratings of sociality, valence, and arousal, and experimenter-coded agent counts (0-3 persons). We correlated individual behavioral RDMs (assessed in Experiment 1) with stimulus feature RDMs (assessed in online pre-study) and tested Kendall’s *τ_a_* values against chance using sign permutation tests (5000 iterations). Multiple comparisons were corrected for family-wise error (FWE) using the maximum statistic approach ^68^. The noise ceiling was calculated by correlating each participant’s RDM with the averaged RDM computed from all participants (upper bound) and with a leave-one-out averaged RDM excluding that participant (lower bound) ^69^.

To quantify the unique and shared contributions of visual, action-related, and affective-communicative features, we performed cross-validated variance partitioning separately for younger and older adults ^2,70,71^. We focused on the eight features that showed significant correlations with behavioral data in both younger and older adults: visual (environment, i.e., indoor/outdoor), action-related (action category, effectors, transitivity, activity), and affective-communicative (valence, arousal, communicativeness). The procedure involved 100 iterations of split-half cross-validation. For each age group, in each iteration, participants were randomly divided into two halves (*n*=22 per half for younger adults, *n*=22 for older adults). The averaged RDM from one half was used to fit seven linear regression models corresponding to all possible combinations of the three stimulus feature sets: (1) visual + action + affective-communicative, (2) action + affective-communicative, (3) visual + affective-communicative, (4) visual + action, (5) visual, (6) action, and (7) affective-communicative. No multicollinearity was detected among the 8 predictors (maximum variance inflation factor = 1.19). We then generated predictions from each model and correlated them with the averaged RDM of the held-out half of participants. The Kendall’s 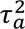 of the correlation quantified the variance explained by each model. Unique variance for each feature set was computed as the difference between full and reduced models (e.g., unique variance explained by visual = Kendall’s 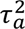 (full) - Kendall’s 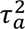 (action + affective-communicative). Shared variances (pairwise and three-way) were derived from intersection of variance explained across models. Variance contributions were tested against chance using one-tailed sign permutation tests (5000 iterations, omnibus-corrected). Differences in accounted variance between feature sets were examined with two-sided Wilcoxon signed-rank tests.

#### MEG Data Preprocessing

MEG data offline preprocessing followed the FLUX pipeline ^72^ using MNE-Python ^73^. First, the Signal-Space Separation (SSS) was applied to reduce environmental artifacts ^74^. Muscle artifacts were identified by bandpass filtering magnetometer data (110-140 Hz) and z-scoring. Segments exceeding subject-specific z-score thresholds were annotated for removal. To further correct for ocular and cardiac artifacts, we applied Independent Component Analysis (ICA). Components containing cardiac artifacts and eye blinks were identified based on their time courses and topographies, resulting in the removal of 2 to 5 independent components per participant. The processed data were then segmented into epochs (−200 to 1000 ms relative to stimulus onset). Catch trials were excluded, leaving 2040 trials per participant. We then removed trials containing erroneous button presses (0.26% of all trials from all participants). Finally, we manually inspected all trials to identify and reject those with excessive noise or high variance. This involved a comprehensive visual inspection using the ft_rejectvisual function in FieldTrip ^75^ and visual check of each individual trial data. On average, 1.40% of the trials (range: 0.05-5.25%) were rejected due to artifacts (see Supplementary Fig. S12 for evoked responses).

#### Time-resolved RSA of MEG data

To investigate the brain temporal dynamics of feature processing during action perception, we performed time-resolved RSA on the MEG data. Neural representational dissimilarity matrices (RDMs) were computed using the MNE-RSA toolbox ^76^. The MEG data were downsampled to 200 Hz. A sliding temporal window of 10 ms radius was used and only gradiometer channels were included. Pairwise Euclidean distances between all MEG data of the associated stimuli pairs were computed to construct the neural RDM.

We first investigated whether MEG activity patterns reflected perceived action similarity. To do this, we correlated neural RDMs (Experiment 2) with a common behavioral RDM obtained from the multi-arrangement task (Experiment 1) averaged across all participants. The resulting Kendall’s *τ_a_* values were tested against chance using one-tailed sign permutation test separately for each condition and each group. Multiple comparisons across time were corrected using a cluster-based permutation test ^77^. The cluster-forming threshold was set to *α* = 0.05, and the cluster-level significance was determined by the distribution of the maximum cluster sums (*α* = 0.05; 5,000 permutations). We further examined the relation between time-resolved neural RDMs and the stimulus features that drove similarity judgments from Experiment 1. For each stimulus feature, the same time-resolved RSA procedure and statistical testing steps as described above for the behavior-neural correlations were applied to the neural-feature correlations.

To further quantify the unique contribution of visual, action-related and affective-communicative features to neural activity patterns, we also performed cross-validated variance partitioning analysis on the time-resolved neural RDMs separately for each age group and condition. In each of the 100 iterations, participants were randomly divided into two groups, following the same cross-validation procedure used for the multi-arrangement data obtained in experiment 1. Specifically, we fitted linear models on one half of the participants and tested predictions on the held-out half, using Kendall’s 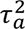 as the prediction metric. The analysis included the 8 feature RDMs that were behaviorally relevant, grouped into three feature sets: visual, action-related and affective-communicative features. In addition to these behaviorally relevant features, the GIST model was included as a measure of low-level visual features that have been shown to account for brain responses in early visual cortex ^4^. We fitted seven different model combinations of three feature sets using the same procedure as done in Experiment 1. Statistical significance was assessed using sign-permutation tests (5,000 iterations) with cluster-based correction for multiple comparisons (cluster-forming threshold *p* < 0.05, cluster-level *α* = 0.05).

To examine the temporal dynamics of feature processing, we conducted a series of comparisons on the effect onset distributions across the 100 split-half iterations. For the original (unblurred) viewing condition, we assessed the temporal sequence of the three feature sets (visual, action-related, affective-communicative) within each age group, and compared effect onsets between age groups. The same analyses were conducted for the blurred viewing condition. Finally, to assess how blurring affected feature processing differently between age groups, we tested main effects of group and condition and their interaction using permutation-based tests. Wilcoxon signed-rank tests were applied to paired comparisons, such as temporal-sequence analyses within a condition and condition comparisons within a group, while Mann-Whitney U tests were used for between-group comparisons. These tests are appropriate for onset distributions derived from permutation-based analyses ^2^.

#### MEG source reconstruction

Individual T1-weighted MRI structural scans were acquired using a 3T Siemens Prisma scanner with 32- or 64-channel head coils and processed with FreeSurfer ^78^ for cortical reconstruction. Source reconstruction was performed using MNE-Python. A single-shell boundary elements model was constructed using the inner skull surface obtained from FreeSurfer. To construct the forward model, the MRI images were manually coregistered with the MEG sensor coordinates with head digitization points. For two participants without MRI structural scan, the fsaverage template was warped to match their digitized head shape. Surface-based source spaces were defined on the cortical surfaces with octahedral downsampling (oct5). Then a surface-based forward model was generated, in which the lead field matrix was calculated based on the source positions relative to the MEG sensor array. A noise covariance matrix was estimated from baseline period (−0.2 s to 0 s before the stimulus onset) of the epoched data. The inverse operator was then constructed using minimum norm estimation with dynamic statistical parametric mapping (dSPM) normalization ^79^. Source estimates were computed by applying the inverse operator to the epoched data (see preprocessing section). Individual source estimates were morphed to the fsaverage template using surface-based spherical registration, ensuring vertexwise cortical anatomy correspondence across participants. On the fsaverage template, the oct5 source space comprised 1,026 vertices per hemisphere.

#### Searchlight RSA on MEG source estimates

For computational efficiency, data were downsampled to 100 Hz for source-level analyses. Spatiotemporal searchlights (spatial radius: 2 cm; temporal radius: 40 ms) were defined at each vertex, within which pairwise Euclidean distances were computed to generate neural RDMs using the MNE-RSA toolbox. Next, we estimated the unique contribution of each feature set using the same split-half cross validation procedure (100 iterations) as employed in the time-resolved RSA section for each searchlight patch. For statistical evaluation, we focused on the age-common time intervals identified in the time-resolved analyses that were significant for both younger and older adults under the original viewing condition. This approach ensured that source-level results were restricted to periods in which both age groups showed reliable feature-specific effects. The same time period was applied to the blurred condition to ensure comparability across conditions. Within each defined time window, Kendall’s 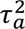 values were averaged. Finally, to test whether the unique variance explained by each feature set was significantly greater than zero, we performed a one-tailed sign-flipping permutation test separately for each age group and viewing condition. Multiple comparisons across patches were corrected using the max-statistic approach ^68^ (10,000 permutations, FWE *p* < 0.05).

## Data availability

The behavioral and preprocessed MEG data and results are available in an Open Science Framework repository (https://osf.io/tk7jf/). Source data are provided with this paper.

## Code availability

Analysis code is available on the same Open Science Framework repository.

## Author contributions

X.-R.P. and S.-C.L. conceived the study with input from C.F.D., Q.H., and A.L. X.-R.P. programmed the tasks and collected the data. X.-R.P. analyzed the data with assistance from Q.H. and input from S.-C.L., A.L. and C.F.D. X.-R.P. wrote the initial draft of the manuscript. S.-C.L., A.L., Q.H. and C.F.D. provided comments for revising later versions of the manuscript. S.-C.L. and C.F.D. acquired the funding.

## Acknowledgements

We gratefully acknowledge the invaluable assistance of Yvonne Wolf-Rosier for MEG data collection, Burkhard Maess for support with MEG experimental implementation, Kerstin Schumer for extensive administrative support, Elisabeth Murzik for participant recruitment, Julia Hofschildt, and Aisuluu Berdalieva for participant recruitment and data collection. X.-R.P. is supported by a grant from the China Scholarship Council (CSC No. 202006990013). S.-C.L. is supported by a grant from the German Research Foundation (DFG, Deutsche Forschungsgemeinschaft) as part of Germany’s Excellence Strategy – EXC 2050/1 & EXC 2050/2 – Project ID 390696704 – Cluster of Excellence “Centre for Tactile Internet with Human-in-the-Loop” (CeTI) of Technische Universität Dresden). X.-R.P. is also partially supported by this funding.

