## Supplementary Materials for "Aging and reorganization of the neural hierarchy underlying action perception: older adults prioritize affective-communicative information"

### Text S1: Pre-study of stimulus feature ratings

An independent sample of participants provided ratings for all the photo stimuli used in the study to characterize their multidimensional features. While basic visual features can be extracted algorithmically or labeled manually, action-related and affective-communicative features are more abstract and subjective, thus require behavioral ratings. These ratings were used to construct feature-specific representational dissimilarity matrices (RDMs) for investigating the organizing principles of observed daily actions and to examine potential age-related differences in feature perception.

**Participants.** A total of 125 participants (62 younger adults and 63 older adults) completed the online rating task via SoSci Survey (https://www.soscisurvey.de/). One younger adult was excluded due to response invariance (using only two of five possible rating values). The final sample comprised 61 younger adults (age range: 18–30 years; M ± SD: 22.87 ± 3.45 years; 28 male, 33 female) and 63 older adults (age range: 63–80 years; M ± SD: 71.95 ± 4.60 years; 39 male, 24 female). All participants gave their informed consent online, the study was approved by the ethics committee of the University of Leipzig (341/23-ek).

**Procedure.** The 102 photographs were divided into three sets of 34 images, with each set rated by different groups of younger and older participants. Participants rated six features on 5-point scales: sociality (1 = not at all social, 5 = very social), valence (1 = very unpleasant, 5 = very pleasant), arousal (1 = very calm, 5 = very intense), activity (1 = no action, 5 = very active), communicativeness (1 = not communicative, 5 = communicative; reflecting the extent to which actions convey social messages or intentions), and transitivity (1 = not at all, 5 = very much; reflecting person-object interactions). The task was self-paced with no time constraints, and participants could pause and resume at any time.

**Interindividual consistency**. To assess the interindividual consistency of feature ratings, we conducted a leave-one-out analysis to calculate Kendall's correlation between each participant's ratings and the mean ratings of all other participants within their age group. This analysis was performed separately for each of the six dimensions (transitivity, activity, arousal, valence, communicativeness, and sociality) and each age group (younger and older adults). Overall, the ratings showed good internal consistency across all dimensions in both age groups, with mean Kendall's tau values ranging from 0.60 to 0.80 (see Fig. S1). Mann-Whitney U tests revealed significant age group differences in two dimensions: communicativeness ($Z = 3.93$, $p < 0.001$) and Sociality ($Z = 2.37$, $p = 0.026$), with older adults showed lower inter-subject agreement than younger adults. No significant age differences were observed for Transitivity ($Z = 0.04$, $p = 0.962$), Activity ($Z = 1.10$, $p = 0.272$), Arousal ($Z = -0.48$, $p = 0.631$), or Valence ($Z = -1.55$, $p = 0.122$). These results indicate that while both age groups showed good consistency in their ratings overall, younger adults showed greater consensus specifically when evaluating the social and communicative aspects of the observed actions.

**Age differences in feature ratings.** To examine potential age-related differences in feature perception, we conducted analyses with linear mixed-effects models (LMMs) for each of the six dimensions, with age group as the fixed effect and random intercepts for participants and stimuli. LMMs were fitted using the lme4 package (Bates et al. 2015), with *p*-values obtained via lmerTest package (Kuznetsova, Brockhoff, and Christensen 2017) using Satterthwaite's approximation for degrees of freedom. Estimated marginal means were calculated using the emmeans package (Lenth 2020). Results showed significant age group differences in three dimensions (see Fig. S1): Older adults provided higher ratings than younger adults for communicativeness ($F_{1,\ 120.03} = 15.76$, $p < 0.001$), sociality ($F_{1,\ 120.05} = 64.85$, $p < 0.001$), and Valence ($F_{1,\ 119.76} = 77.68$, $p < 0.001$;). No significant age differences were observed for Arousal ($p = 0.597$), Activity ($p = 0.517$), or Transitivity ($p = 0.530$). These findings suggest that older adults tend to perceive the observed daily actions more as communicative and more pleasant compared to younger adults, while both age groups show similar perceptions of the actions' arousal level, activity level, and degree of object interaction.

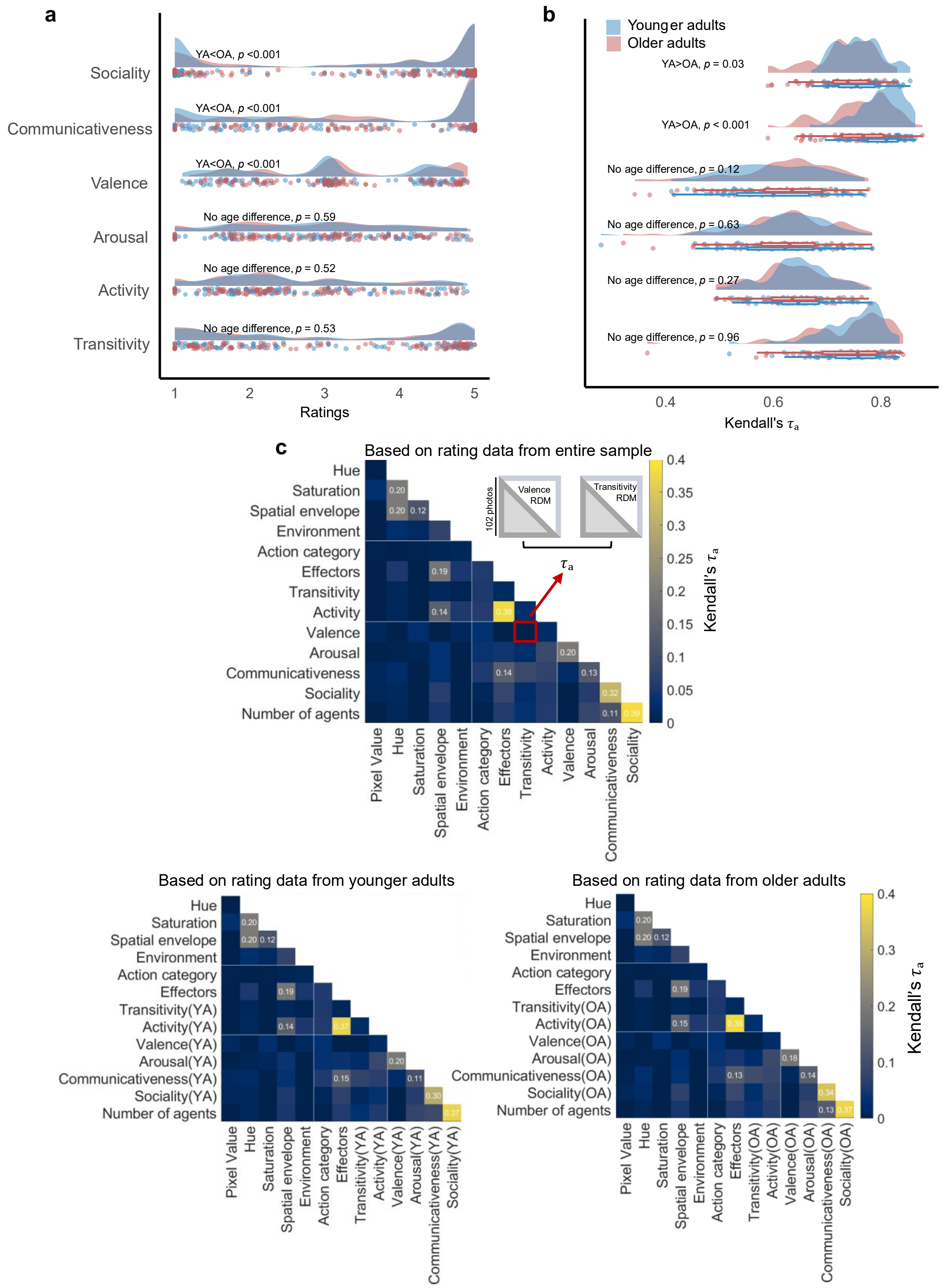


**Fig. S1** Stimulus feature space characterization. **a** Distribution of raw ratings (1-5 scale) for each dimension by age group. **b** Leave-one-out inter-individual consistency analysis showing correlations (Kendall's $\tau_a$) between each participant's ratings and the mean ratings of others in their age group. **c** Between-feature correlation matrix showing Kendall's $\tau_a$ correlations (based on rating data from entire sample).

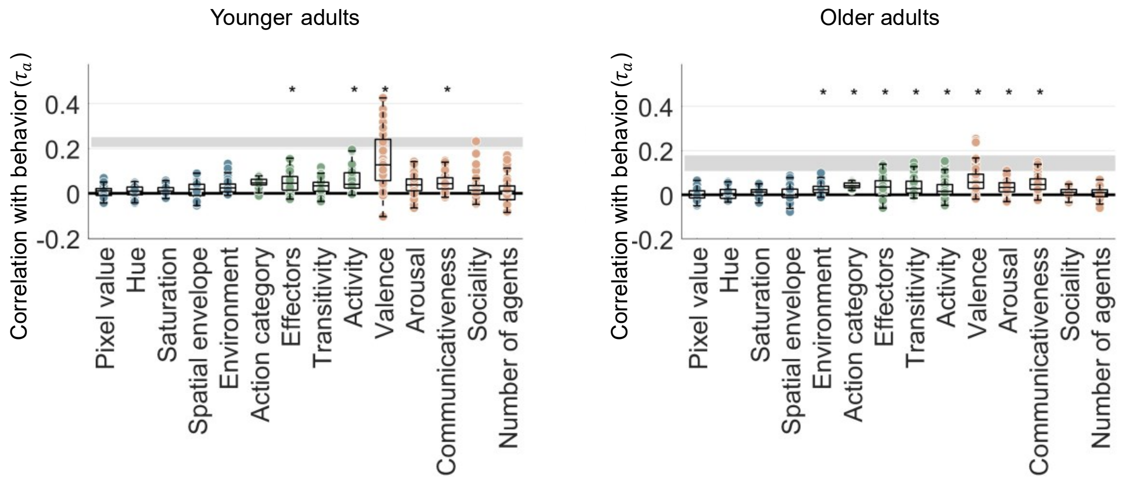


**Fig. S2** Correlations between participants' behavioral representational dissimilarity matrices (RDMs) and stimulus feature RDMs in younger and older adults (Experiment 1). Each dot represents one participant (values of individual participants overlap). Asterisks indicate significant correlations (FWE-corrected $p < 0.05$, sign-permutation tests).

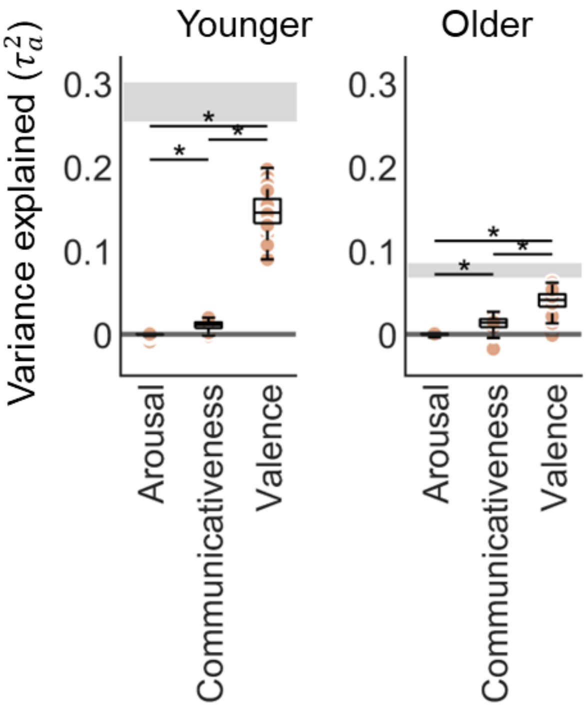


**Fig. S3** Unique variance explained by the communicativeness, valence, and arousal, analyzed separately for younger and older adults using cross-validated half-split variance partitioning in the behavioral data from Experiment 2. Each dot represents one iteration from 100 half-split cross-validations. Asterisks indicate significant differences between feature sets (all $p < 0.001$, Wilcoxon signed-rank tests). This control analysis suggests that valence explain most of the variance in the data in both age groups, beyond the arousal and communicativeness. Notably, in younger adults, only valence showed significant unique contribution above zero ($p < 0.001$), whereas in older adults both communicativeness ($p = 0.002$) and valence ($p < 0.001$) showed significant unique contributions; arousal did not yield significant unique contribution in either group.

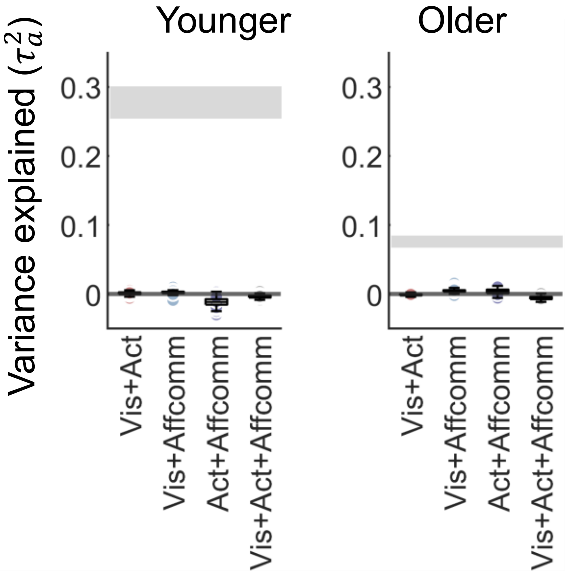


**Fig. S4** Shared variance explained by the visual (vis), action (act), and affective-communicative (affcom), analyzed separately for younger and older adults using cross-validated half-split variance partitioning in the behavioral data from Experiment 2. Each dot represents one iteration from 100 half-split cross-validations. All shared variances among feature sets did not reach significance (*p*s > 0.20).

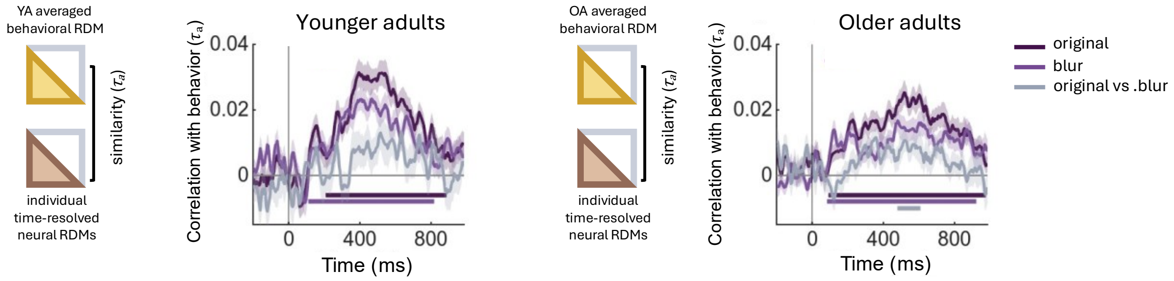


**Fig. S5 Control analysis of neural activity track age-specific perceived action similarity over time**. Time course of correlations between age-specific behavioural RDMs and neural RDMs are shown for younger adults and older adults in two viewing conditions. Age group specific behavioral RDMs were calculated by separately averaging data from Experiment 1 for each of the age group first. The age-specific behavioral RDM was then correlated with individual time-resolved MEG RDMs from Experiment 2. Significant time points (sign-permutation testing, cluster-corrected $p < 0.05$) are indicated with horizontal lines below the curves (along the x-axis). Shaded areas represent standard error of the means across participants. The overall pattern of results is consistent with the main analysis using the grand-averaged behavioral RDM computed across both age groups (Fig. 3 in the manuscript), i.e., no significant clusters were observed between brain-behavior correlations in younger adults between the two viewing conditions (i.e., original vs. blur), whereas in older adults a significant cluster was observed around 491-607 ms after stimulus onset.

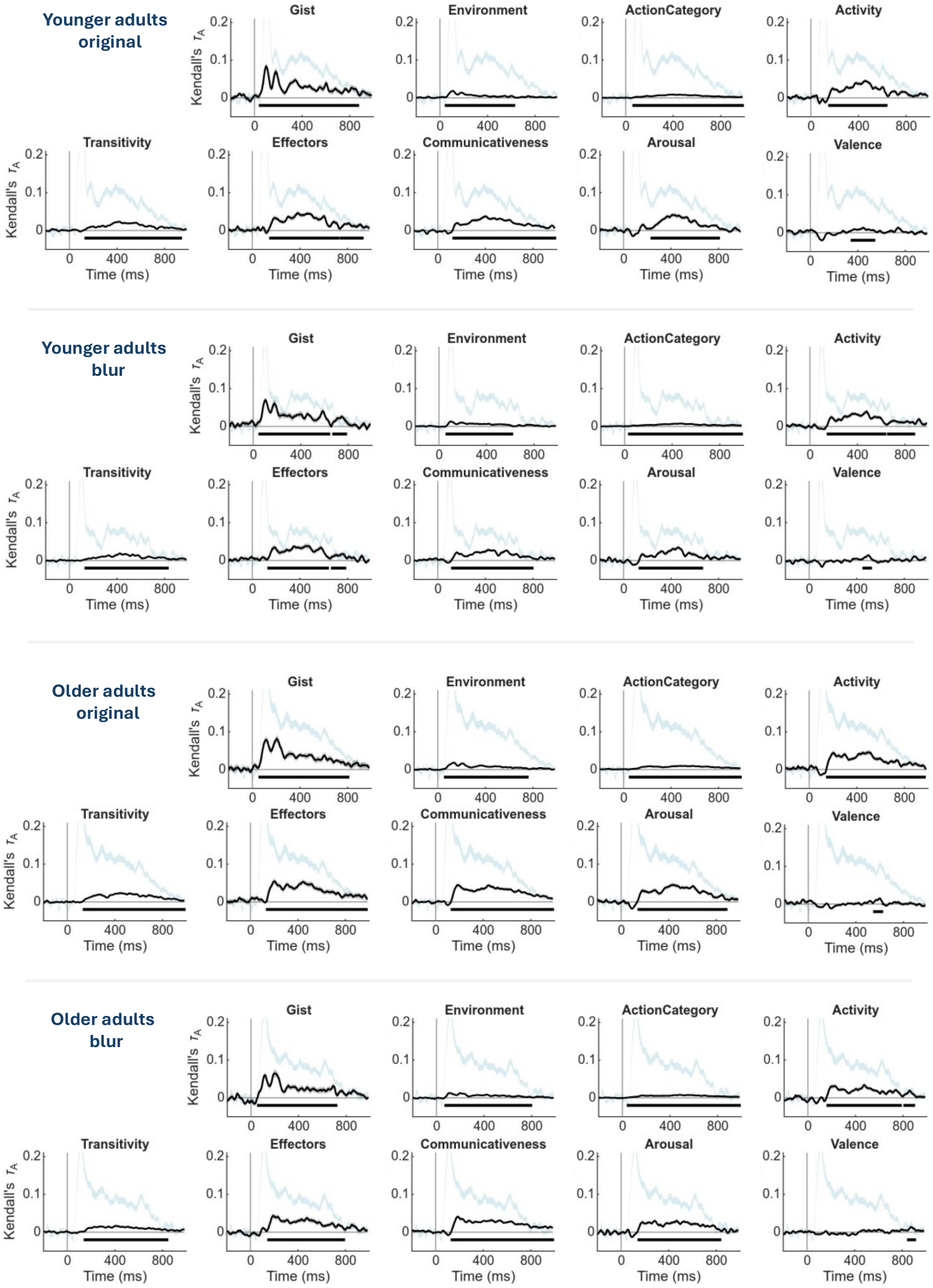


**Fig. S6 Correlations between stimulus features and the time-resolved neural representational dissimilarity matrices (RDMs) in two age groups and viewing conditions.** Significant time points (sign-permutation testing, cluster-corrected $p < 0.05$) are indicated with horizontal lines below the curves (along the x-axis). The noise ceiling is shown in light blue (leave-one-subject-out correlation, mean ± SEM).

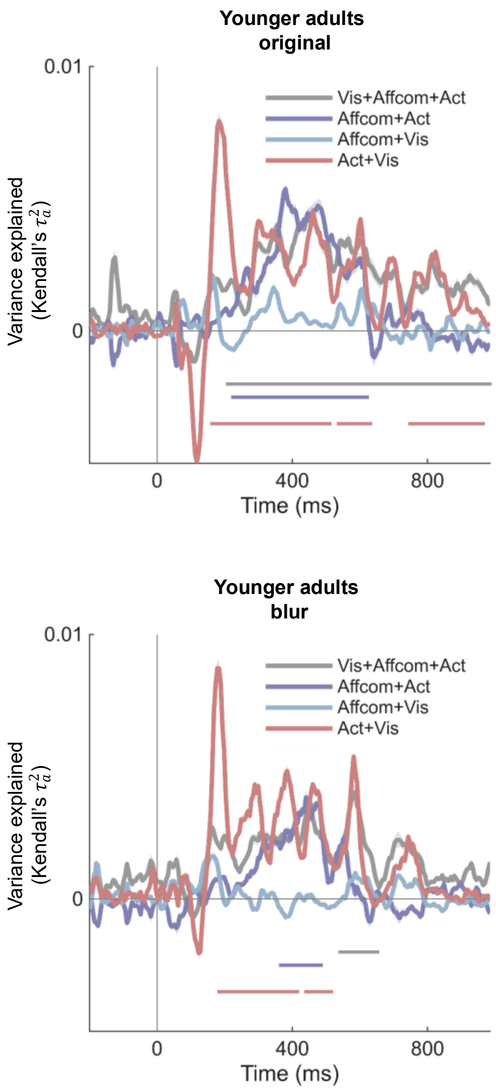


**Fig. S7** Time course of shared variance among visual (vis), action (act), and affective-communicative (affcom) predictors in the cross-validated variance partitioning analysis for the two age groups and viewing conditions. Significant time points (sign-permutation testing, cluster-corrected $p < 0.05$) are indicated with horizontal lines below the curves (along the x-axis).

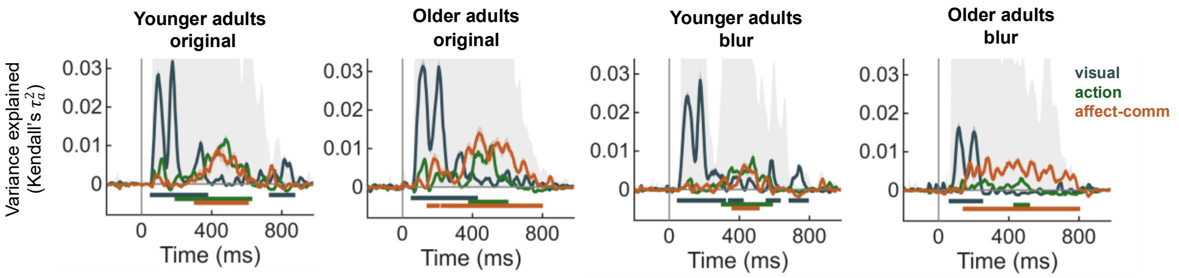


**Fig. S8 Control analysis of neural processing dynamics of unique visual, action and affective-communicative features using age-group-specific feature ratings.** Replicates the analysis shown in Fig. 4b in the manuscript, but feature RDMs for transitivity, activity, communicativeness, valence, and arousal were constructed using ratings collected from the corresponding age group (i.e., age-group specific ratings). Shaded areas indicate standard error; gray shaded backgrounds indicate the noise ceiling. Significant time points (sign-permutation testing, cluster-corrected $p < 0.05$) are marked with horizontal lines along the x-axis. The results show a pattern as observed in the main analysis reported in the manuscript, confirming that the findings are robust regardless of whether pooled or age-specific ratings are used.

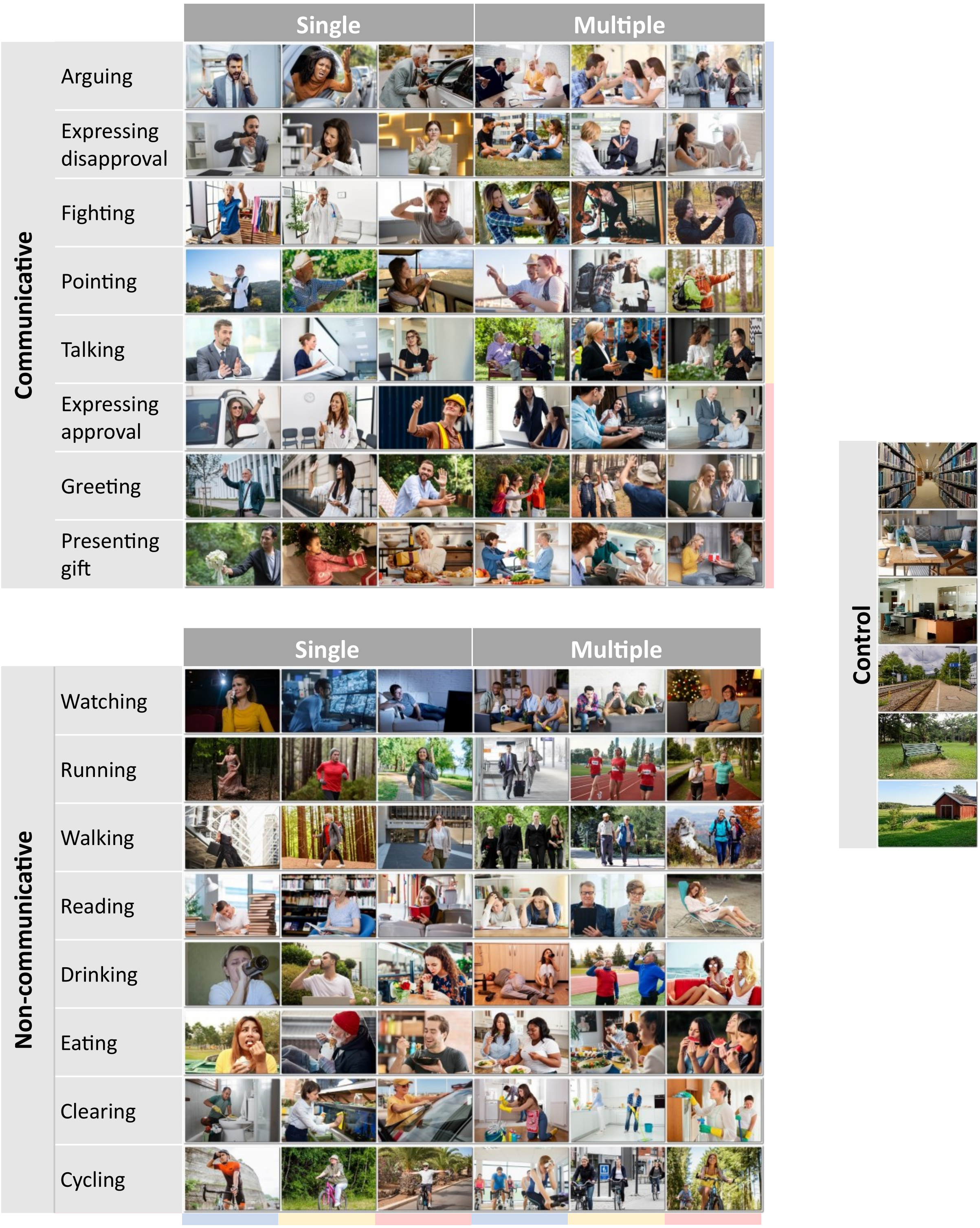


**Fig. S9 Overview of the experimental stimulus set.** The stimulus set comprises 96 action photos depicting 8 communicative actions and 8 non-communicative actions. Each action is represented by 6 photographs: 3 with single agents and 3 with multiple agents. Six control photos depicting scenes without actions or agents are also included. All images were originally purchased from Shutterstock (https://www.shutterstock.com) and are reproduced with permission under license from Shutterstock.

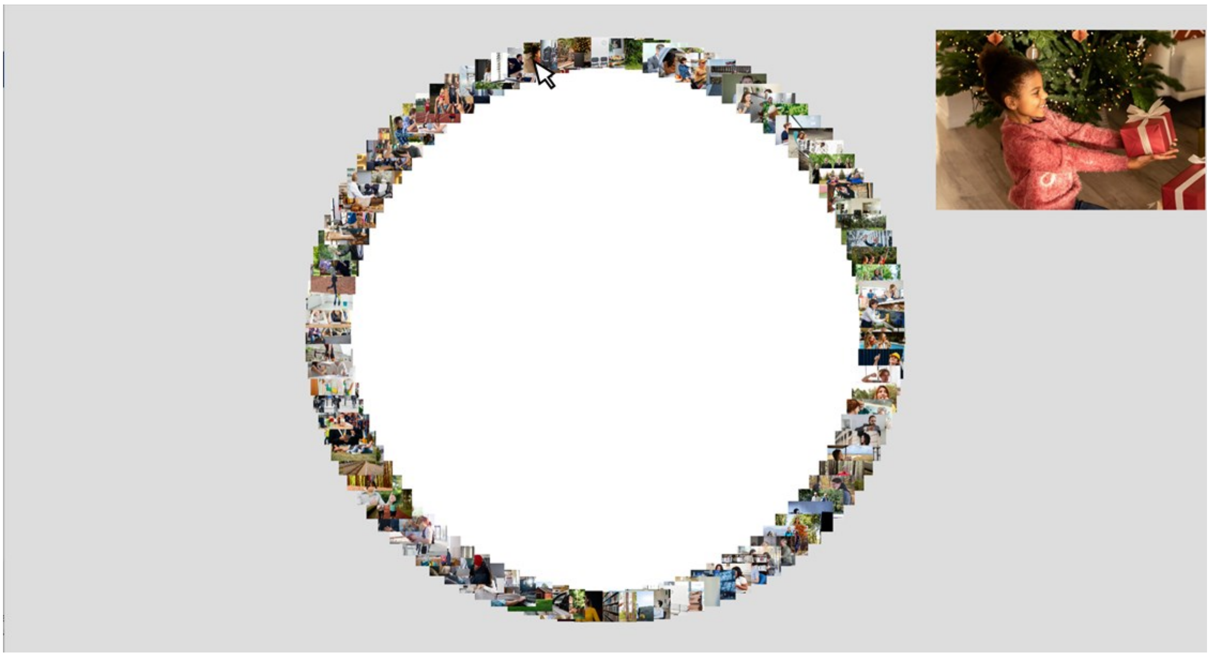

**Fig. S10 Computer interface of the multi-arrangement task (state at the start of the initial trial is shown).** In the first phase of the task ("arrange all items"), participants were presented with all 102 action stimuli arranged in a circular arena. They were instructed to drag and drop items to reflect perceived similarity. To accommodate all items simultaneously on the screen, stimuli were initially displayed as small thumbnails. To facilitate the inspection of visual details, a larger preview of the image appeared dynamically when the participant hovered the mouse cursor over a thumbnail (as illustrated by the enlarged photo in the top right corner). Images are reproduced under license from Shutterstock (https://www.shutterstock.com/).

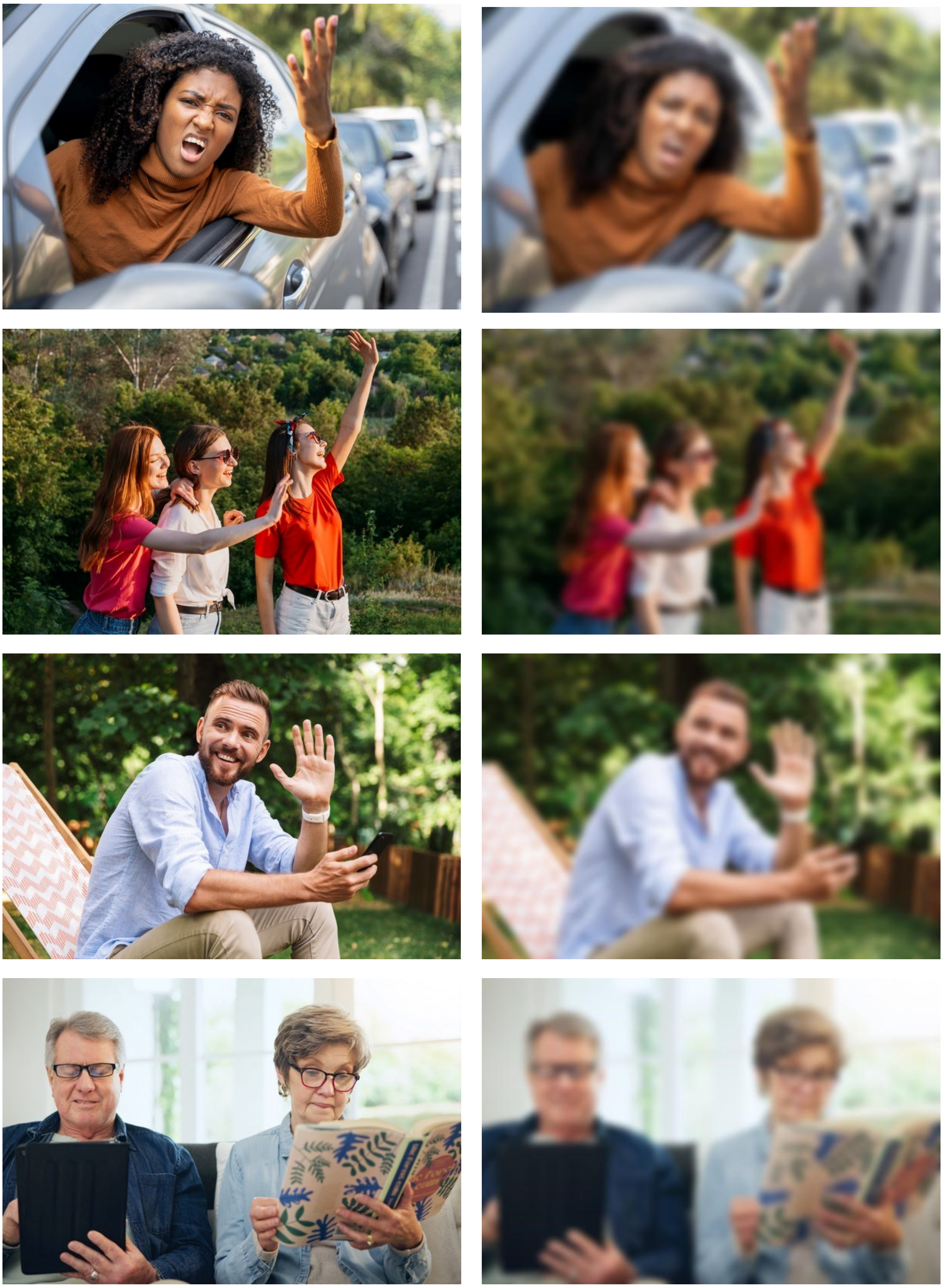

**Fig. S11 Example images in original (unblurred) and blurred versions.** Left, original; right, blurred. Images are reproduced under license from Shutterstock (https://www.shutterstock.com/).

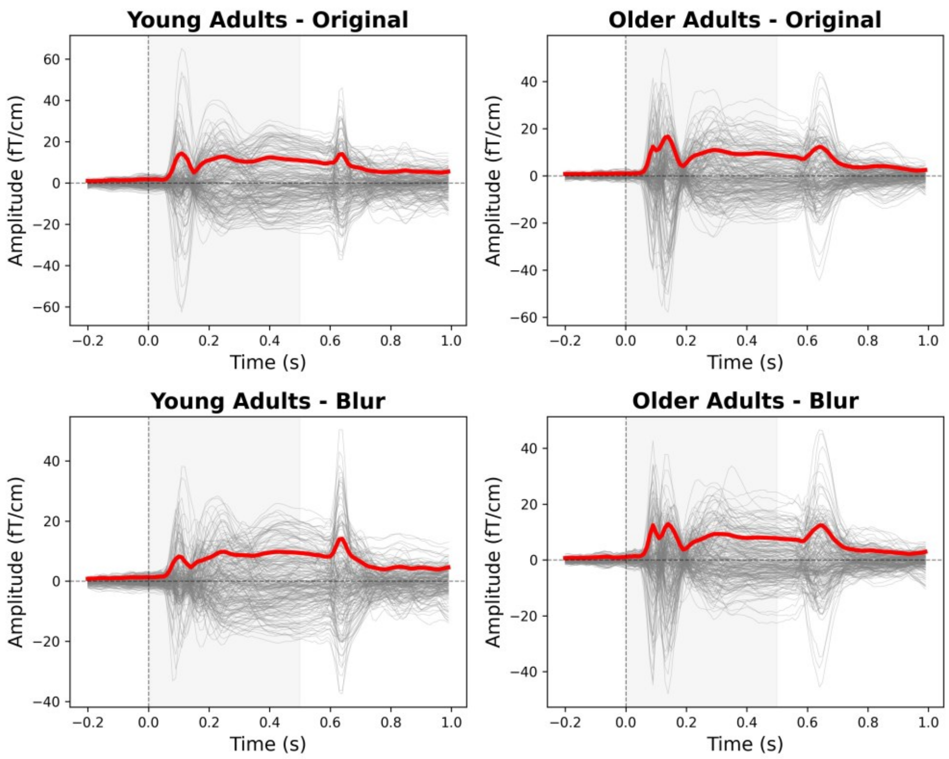


**b**

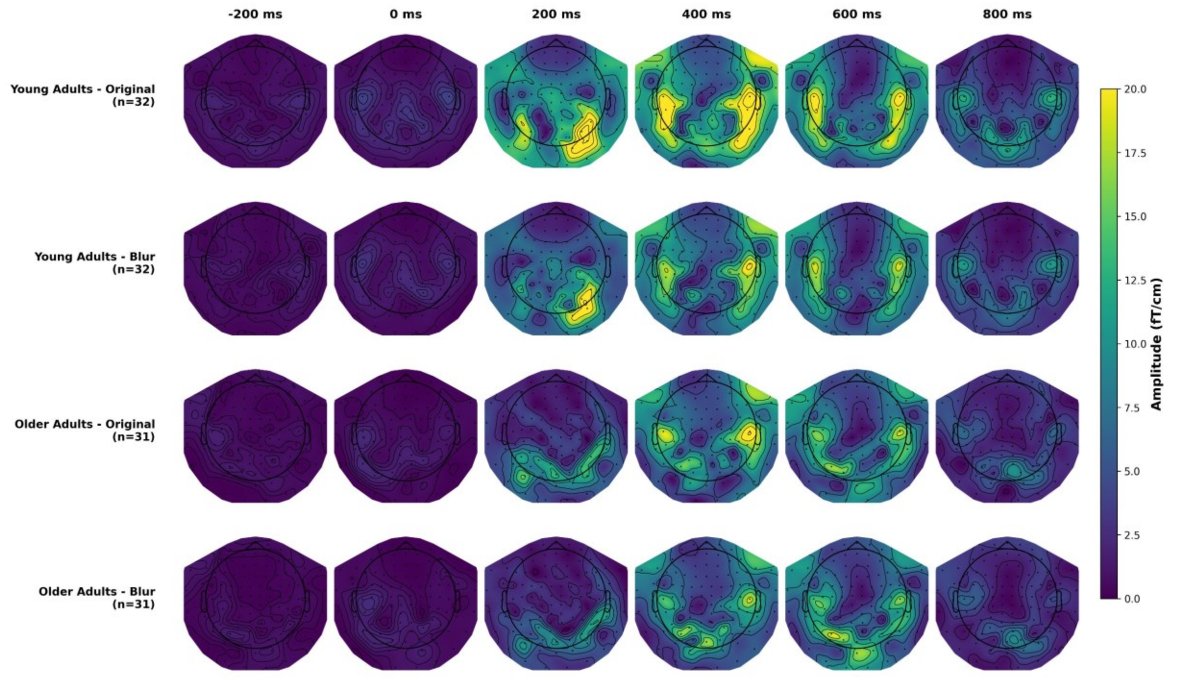


**Fig. S12 Global field power and topographic distribution of evoked magnetic responses. a** Gray lines represent individual gradiometer channels (n = 204 channels). Red lines indicate the Global Field Power (GFP), calculated as the standard deviation across all channels at each time point. GFP provides a reference-independent measure of overall response strength, avoiding potential biases from channel averaging (Lehmann and Skrandies 1980). The gray shaded area (0–0.5 s) marks the stimulus presentation period. Vertical dashed line indicates stimulus onset. **b** Topographic distribution across time. Spatial distribution of evoked responses at six time points (-200 to 800 ms).

Table S1 Demographic and cognitive characteristics of participants in the MEG experiment

| | YA<br>*M (SD)* | OA<br>*M (SD)* | YA vs. OA<br>*t (p)* |
|---|---|---|---|
| *N* | 32 | 31 | - |
| Age | 25.16 (3.63) | 71.58(3.44) | -51.20 (< 0.001) |
| Gender | 15M / 17F | 17M / 14F | 0.00[a] (1) |
| MoCA | - | 27.48(1.90) | - |
| Duke Social Support Index | | | |
| Social interaction | 9.01 (1.00) | 8.90 (1.44) | 0.28 (0.78) |
| Social support | 20.43 (0.79) | 20.29 (1.01) | 0.57 (0.58) |
| Visual search task | | | |
| Accuracy [%] | 96.02 (3.23) | 94.23 (4.46） | 1.82 (0.07) |
| RT [ms] | 1501.34 (231.86) | 2417.21 (473.65) | -9.80 (<0.001) |
| Identical-Pictures Test | | | |
| Score | 33.88 (4.06) | 21.74 (3.30) | 12.99 (<0.001) |
| RT [ms] | 2052.64 (292.55) | 3341.43 (642.99) | -10.29 (<0.001) |
| Spot-the-Word Test | | | |
| Accuracy [%] | 67.84 (8.12) | 80.24 (9.24) | -5.67 (<0.001) |
| RT [ms] | 4981.96 (1380.13) | 5821.55 (2422.44) | -1.70 (0.09) |

*Note.* YA, younger adults; OA, older adults; M, mean; SD, standard deviation. [a]$\chi^2$ value. Cognitive measures included the Spot-the-Word Test (verbal knowledge), Identical-Pictures Test (processing speed), and a visual search task (visual attention). Social support was assessed using the Duke Social Support Index. See Methods for further details.

**Appendix: English translations of the instruction of the multiple arrangement task (Experiment 1).**

How do we perceive, interpret, and understand different actions of others? This is the intriguing question we aim to answer in this study. In the current task, we will present you with 102 photos of human(s) performing various actions. Your task is to arrange the photos based on your own subjective perception of their similarities. Here's a step-by-step guide for what you need to do in the task:

**Step 1: Get an Overview of the Photos**

Before starting to sort the photos, please take a moment to get an overview of all the photos. Simply hover over each photo arranged along the circle with the mouse; the enlarged version of each photo will then appear on the right side of the screen. This will help you form an initial impression of the spectrum of actions depicted in the photos.

**Step 2: Sort the Photos**

After you have had an overview of the photos, the next step is to sort them. To do so, use the mouse to drag and drop each of the photos one at a time and put it in a position within the circular empty space. The similar the photos the closer they should be placed to each other, and the more different they are from each other, the farther apart they should be placed from each other. There's no correct or wrong ways in sorting the photos; you should sort the photos based on your own subjective perception of their similarities (dissimilarities).

**Step 3: Review Your Sorting**

Once you've sorted all 102 photos, take a moment to review your sorting. Remember, your satisfaction with the arrangement is essential for the study. So, take your time to ensure you are happy with your arrangement.

**Step 4: Take a Break**

After you have completed the initial sorting, our system will review it. During this time, please take a 3 to 5 min break to rest and mentally prepare for the next part of the task. For the software system to gain a better representation of your sorting, the system often requires more information. Therefore, based on your initial sorting results, the system will pick some subsets of photos from the 102 photos to ask you to sort them again.

**Step 5: Re-arrange Some Subsets of Photos**

After your break, the system will begin to present you with subsets of photos from your initial arrangement, particularly those whose similarity-based arrangements were not quite clear yet. Your task remains the same: sort the photos based on your own subjective perception, placing similar photos closer together and dissimilar photos farther apart. The photos will be shown one subset a time; some subset will have more photos than others.

Importantly, please use similar sorting principles to arrange the photos as you did for the initial arrangement. A sudden change in the ways you sort the photos may confuse the system.

Once you're satisfied with the arrangement of a given subset, please click the green button located in the bottom left corner of the screen to confirm it.

This process will repeat across multiple runs with different subsets of photos. After each of such runs, the system gathers more information about how you subjectively perceptive the actions shown in the photos. Your task continues until the system has collected sufficient evidence to represent your sortings or will end after 2 hours.
